# Coupling of NOD2 to GIV is Required for Bacterial Sensing

**DOI:** 10.1101/2022.04.26.489574

**Authors:** Gajanan D. Katkar, Mahitha Shree Anandachar, Saptarshi Sinha, Stella-Rita Ibeawuchi, Celia R. Espinoza, Jane Coates, Yashaswat S. Malhotra, Madhubanti Mullick, Vanessa Castillo, Daniella T. Vo, Debashis Sahoo, Pradipta Ghosh

## Abstract

Sensing of pathogens by Nucleotide oligomerization domain (NOD)-like 2 receptor (NOD2) induces a protective inflammatory response that coordinates bacterial clearance. Polymorphisms in NOD2 impair bacterial clearance, leading to chronic gut inflammation in Crohn’s disease (CD) via mechanisms that remain incompletely understood. We identify GIV/Girdin (CCDC88A) as a NOD2-interactor that shapes bacterial sensing-and-signaling in macrophages. Myeloid-specific GIV depletion exacerbated and protracted infectious colitis and abolished the protective effect of muramyl dipeptide (MDP) in both chemical colitis and severe sepsis. In the presence of GIV, macrophages enhance anti-bacterial pathways downstream of NOD2, clear microbes rapidly and concomitantly suppress inflammation. GIV’s actions are mediated via its C-terminus, which directly binds the terminal leucine-rich repeat (LRR#10) of NOD2; binding is augmented by MDP and ATP, precedes receptor oligomerization, and is abolished by the *1007fs* CD-risk variant which lacks LRR#10. Findings illuminate mechanisms that underlie protective NOD2 signaling and loss of function in the major *1007fs* variant.

**In brief:** This work reveals a mechanism by which macrophages use their innate immune sensor, NOD2, to protect the host against overzealous inflammation during bacterial infections, and the consequences of its loss, as occurs in the most important Crohn’s disease-risk variant.

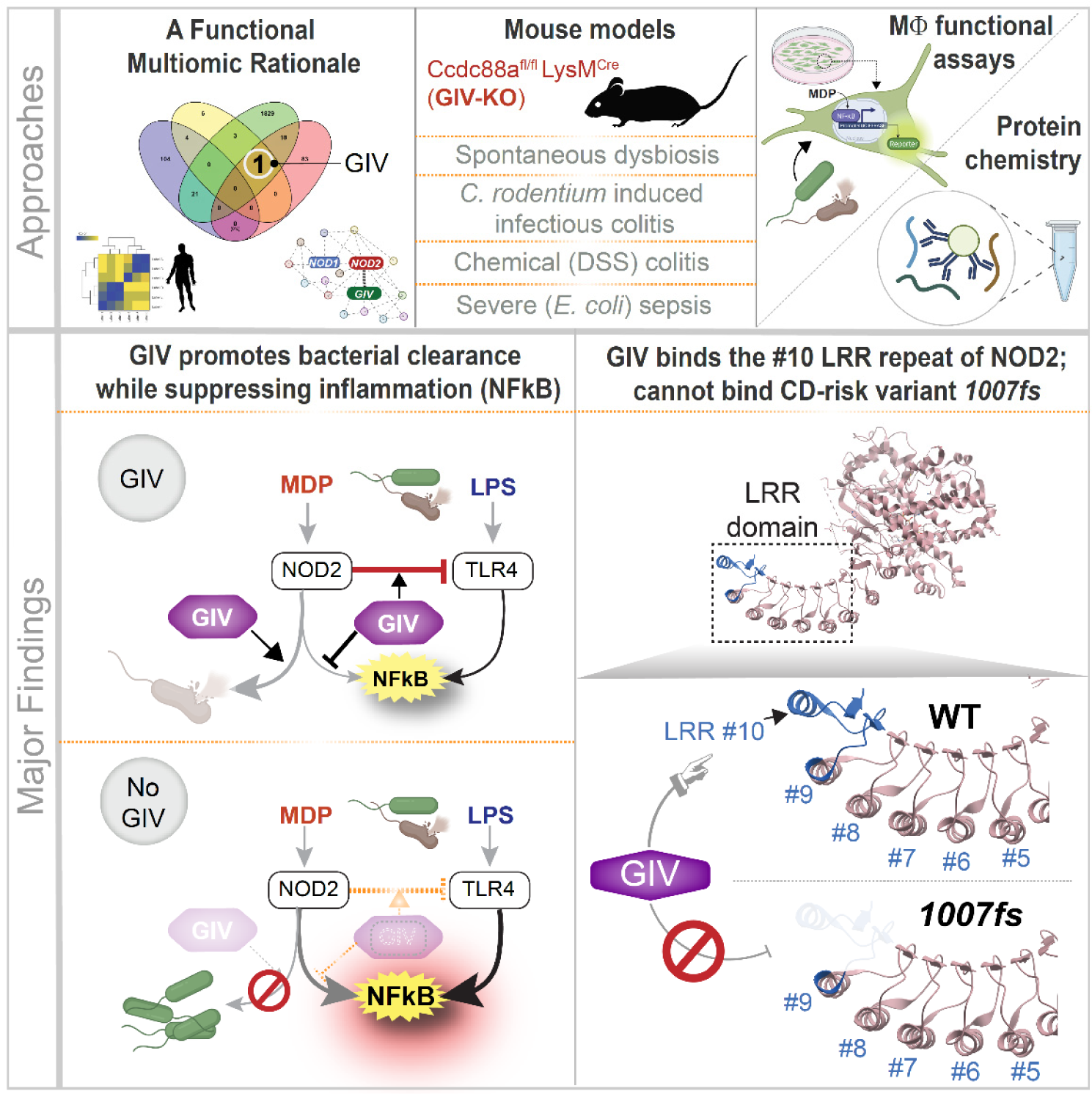
GRAPHIC ABSTRACT

**HIGHLIGHTS:**

- GIV is a functional and direct interactor of the terminal LRR repeat of NOD2
- Mice lacking MФ GIV develop dysbiosis, protracted ileocolitis and sepsis
- MDP/NOD2-dependent protective host responses require GIV
- CD-risk NOD2 *1007fs* variant lacking the terminal LRR#10 cannot bind GIV

## INTRODUCTION

Intestinal macrophages are important for gut development, immunity, homeostasis, and repair (<u>Yang and</u> <u>Cong 2021</u>). Their dysregulation leads to uncontrolled gut inflammation, such as that encountered in Crohn’s disease (CD), a type of chronic gut inflammation within the spectrum of inflammatory bowel diseases (IBD) (Davies and Abreu 2015; Marks 2011).

Among the innate immune pathways that are known to be important, the role of nucleotide binding oligomerization domain containing 2 (NOD2) in microbial sensing in intestinal macrophages is well documented (<u>Marks 2011</u>). Like its counterpart NOD1, NOD2 is a cytosolic bacterial sensor with multifaceted role in innate immune sensing and signaling and plays a vital role in the broad context of host resistance to microbial challenge as well as maintenance of tissue homeostasis (Caruso et al. 2014; Philpott et al. 2014). Sensing muramyl dipeptide (MDP), a constituent of both Gram-positive and Gram-negative bacteria (and a specific ligand for NOD2) and subsequent activation of proinflammatory pathways (e.g., NFκB and MAPK via RIP2 kinase) is a well-characterized function of NOD2 (McCarthy et al. 1998; Nachbur et al. 2015). Such sensing and signaling induces a controlled inflammatory response that coordinates bacterial clearance and confers immunity.

NOD2 was also the first gene identified as a risk factor for ileal CD (<u>Lesage et al. 2002</u>). Several NOD2 variants increase the risk of CD (Kufer et al. 2006; Nelson et al. 2021; Rescigno and Nieuwenhuis 2007; Vignal et al. 2007) and the disease course of ulcerative colitis (UC) (<u>Freire et al. 2014</u>). Of these, the *1007fs* variant displays 100% penetrance and is most consistently associated with CD across multiple studies and population groups (<u>Economou et al. 2004</u>). NOD2 variants are known to confer defective bacterial clearance in macrophages; they also impair NFκB activation (Ashton et al. 2022; Caruso et al. 2014; Graham and Xavier 2020; Warner et al. 2013); however, CD patient’s gut mucosa show increased activation of NF-κB (Eckmann and Karin 2005; Hugot et al. 2001; Maeda et al. 2005; Ogura et al. 2001; Rogler et al. 1998). Based on these observations, NOD2 is believed to restrict activation of the NF-κB pathway by TLR2/4 (Negroni et al. 2018; Watanabe et al. 2014; Watanabe et al. 2008; Watanabe et al. 2004; Watanabe et al. <u>2005</u>) and its dysfunction causes runaway inflammation, thereby increasing the risk of colitis. Despite these insights, mechanism(s) that coordinate NOD2-mediated protective signaling in macrophage, and how/why NOD2 variants such as *1007fs* may be compromised remains poorly understood.

Using trans-scale approaches ranging from computational analyses of the transcriptome, microbiome and proteome, through *in vivo* disease modeling and *in cellulo* interventional studies, to *in vitro* biochemical studies that offer a molecular-level insight, here we reveal that a multi-modular signaling scaffold, GIV (a.k.a Girdin) is a functional and direct NOD2-interactor that shapes innate immune sensing and signaling. We also reveal how NOD2 couples to GIV, and provide a structural basis for defective signaling via the most important CD-risk variant, *1007fs*.

## RESULTS

### Identification of GIV as a putative functional and physical interactor of NOD2

To identify modulators of NOD2-mediated inflammation, in particular, the hyperinflammation that is observed in CD, we overlayed 4 datasets independently generated by 3 groups, each assessing either physical or functional interactions of NOD2 (**Fig 1A**). Only 1 candidate emerged, GIV (a.k.a Girdin; encoded by the *CCDC88A* gene) (circle; **Fig 1A**) that fulfilled 3 key criteria: (i) whose depletion in a genome-wide siRNA screen in HEK cells (<u>Warner et al. 2013</u>) resulted in MDP-induced hyperactivation of NFkB (**Fig 1B**); (ii) which interacts (either directly or indirectly) with NOD2, but not NOD1, as determined by BioID proximity ligation assays in HEK cells (<u>Lu et al. 2019</u>) (**Fig 1C**); and (iii) whose expression tracks NOD2 in the colons from UC/CD-afflicted patients, as determined by RNA seq (<u>Sahoo et al. 2021</u>) (**Fig 1A**). *CCDC88A* is induced in the intestinal macrophages isolated from inflamed UC/CD colons (**Fig 1D**). *CCDC88A* and *NOD2* are co-induced in the UC/CD colon tissues (**Fig 1E**) and reduced in responders to anti-TNFα treatment (**Fig 1E**). The correlation between NOD2 (but not NOD1) and CCDC88A was consistently preserved across numerous independent cohorts (**Fig 1F-G**). Because the multi-modular cytosolic scaffold GIV is known to bind and modulate many cellular sensors, i.e., diverse classes of cell surface receptors (<u>Ghosh and Mullick 2021</u>) and was recently shown to also modulate macrophage responses (<u>Swanson et al. 2020</u>), these findings suggest that this scaffold may also modulate the cytosolic sensor, NOD2.

**Figure 1.**
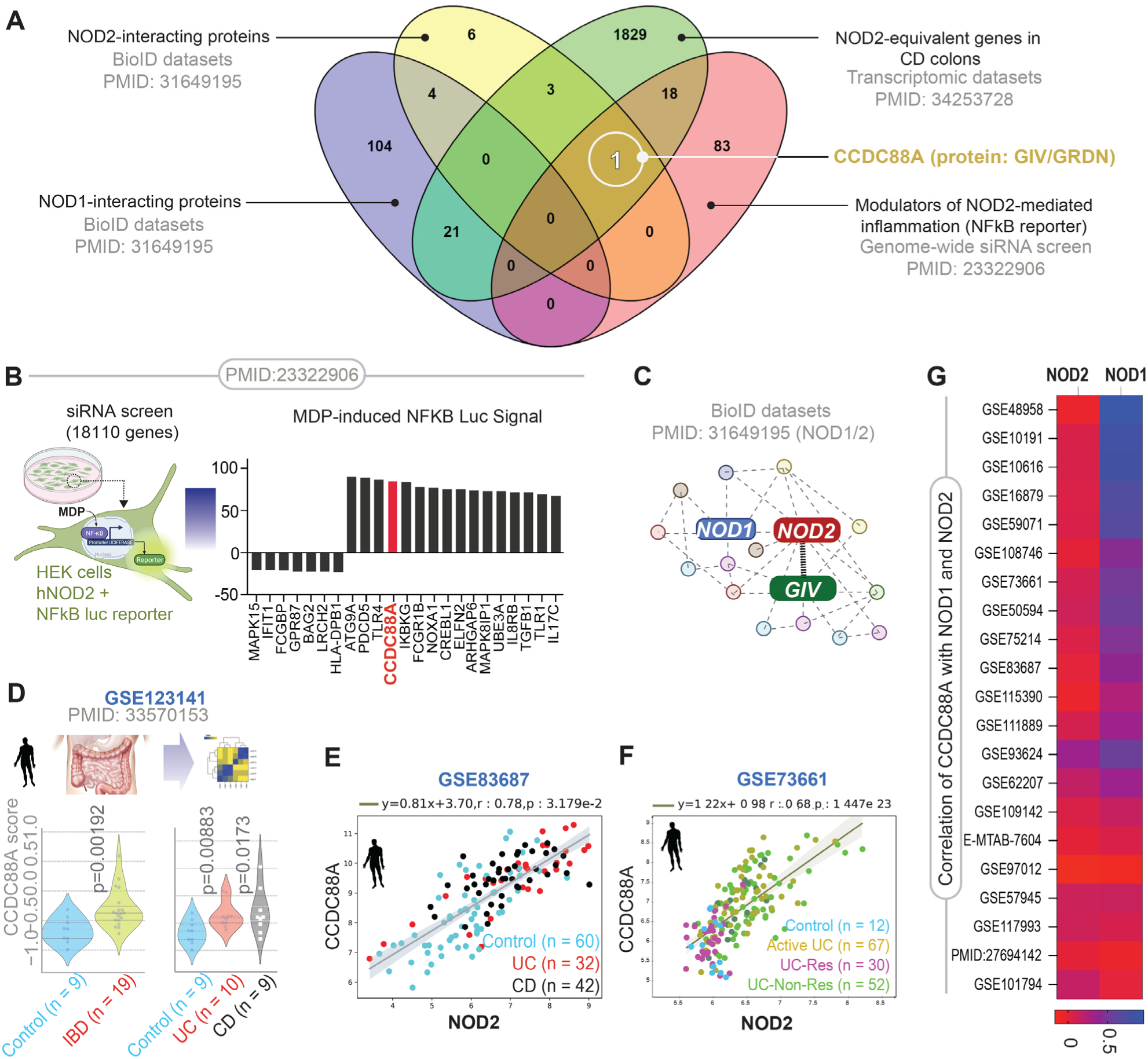
Study premise: Identification of GIV/Girdin as a putative modulator of NOD2 signaling. **A.** Venn diagram showing the number of genes/proteins, identified by independent studies, to associate with NOD1/2 functionally or physically. One gene/protein (CCDC88A/Girdin; white circle) interacts with NOD2 but not NOD1, is upregulated in CD and modulates NOD2-dependent NFkB activation. **B.** Schematic and bar graph display the results of a siRNA-based screen used to identify modulators of NOD2-dependent NFkB activation. Red bars highlight the impact of GIV (CCDC88A)-depletion compared to other prominent players. **C.** Schematic summarizes the findings of BioID proximity labeling studies (<u>Lu et al. 2019</u>) conducted in HEK cells to establish NOD1/2 interactomes. **D.** Violin display the abundance of NOD2 and GIV transcripts in intestinal macrophages. *p* values were determined (compared to control) by Welch’s t-test. **E-F**. Scatter plots of the abundance of CCDC88A and NOD2 transcripts in full thickness colonic tissues from control and IBD subjects, untreated (E) or after treatment (F) with anti-TNFα (Res, responder; Non-Res, non-responder). **G**. Heatmap of correlation coeff. between NOD1/2 and CCDC88A across numerous independent IBD cohorts. *p* value < 0.05 is consider as significant.

### GIV-KO mice develop dysbiosis, and exacerbated and protracted *Citrobacter*-induced colitis

To study the role of GIV *in vivo*, we used a myeloid specific GIV-KO (Ccdc88a^fl/fl^/LysM^Cre^) model (see *Methods*) (<u>Swanson et al. 2020</u>). We began by investigating if the gut microbes in the GIV-KO mice was altered by analyzing fecal pellets of individual mice by 16S RNA sequencing. The alpha and beta diversity indices did not show significant difference between GIV-KO and WT fecal microbiome (**Fig S1**). Surprisingly, despite sharing the same food, water and space with their littermate WT controls, the GIV-KO mice spontaneously develop gut dysbiosis by ∼8-12 wk. All the GIV-KO mice (5/5) had the strain *Rhizobiales* but undetectable in the control littermates (*P* = 0.037) (**Fig 2A-B**). *Rhizobiales* is uniquely present in CD patients, but undetectable in healthy subjects (<u>Kaakoush et al. 2012</u>). Transmission electron microscopy (TEM) confirmed that peritoneal macrophages from GIV-KO mice retain higher number of the CD-associated pathogenic *E. coli* strain (*AIEC* LF82) (**Fig 2C-D**). A classic gentamicin-protection assays were also conducted, which are discussed later in this study.

**Figure 2.**
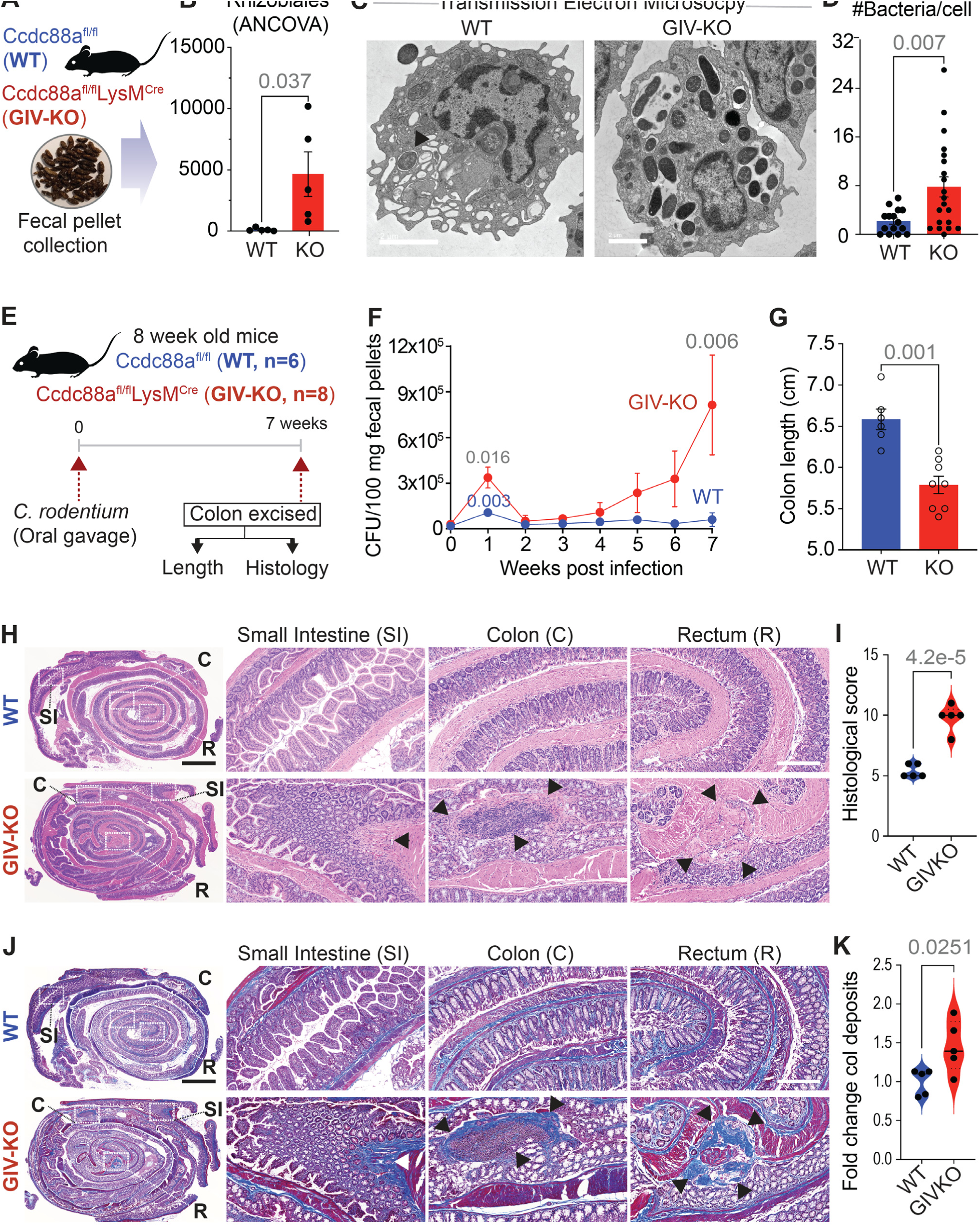
A mouse model of dysbiosis, impaired microbial clearance, patchy chronic transmural ileocolitis and fibrosis. **A-B**. Schematic (A) and bar graph (B) display the process and outcome of a 16S fecal microbiome analysis at baseline in 10 wk-old myeloid-specific (LysMCre) GIV-KO mice and their control littermates (WT). *p* value was estimated by unpaired t-test; n = 5 mice in each group. **C-D**. TEM micrographs (C) display representative images of peritoneal macrophages challenged with live *AIEC* LF82 (MOI 1:30) for 1 h. Scale bar = 2 µM. Bar graphs (D) display findings based on ∼20-30 randomly imaged fields; n = 2 repeats. See also Figure 4E for classic gentamicin-protection assays. **E-K**. Panels describing the experimental design (**E**) and findings (**F-K**) in an infectious colitis model of GIV-KO and control littermates induced using *Citrobacter rodentium* (initially termed *Citrobacter freundii* biotype 4280 (<u>Newman et al. 1999</u>); strain name DBS100; 5 x 10^8^ CFU/200ul/mouse. GIV-KO, n=8; WT, n=6. Findings are representative of two independent repeats. Line graphs (**F**) display the bacterial burden in fecal pellets over a 7 wk period after the initial oral gavage. P values were determined by unpaired t-test. *, < 0.05; **, 0.01. Bar graph (**G**) displays the differences in colon length. *p* values were determined by unpaired t-test. H&E (**H**) or trichrome (**J**)-stained images representative of Swiss rolls of the entire intestinal tract are shown. Scale bar = 2.5 mm. Magnified fields of the rectum (R), colon (C) and small intestine (SI) (on the right) of the corresponding boxed regions (on the left) are shown. Scale bar = 250 µm. Arrows show regions of transmural inflammation/crypt distortion, immune infiltrates (in H) correspond also to transmural fibrosis (in J). Segments in between these patches appear normal. **(I)** Bar graphs show the histology index (<u>Bouladoux et al. 2017</u>) (based on submucosal inflammation, percent area involved, inflammatory infiltrates in LP and crypt hyperplasia and the degree of fibrosis (K), as assessed by H&E and trichrome staining on n = 5 WT and 5 KO mice. *p* values were determined by unpaired t-test. All results are displayed as mean ± SEM.

Next, to study the role of macrophage GIV *in vivo* in maintaining gut microbe homeostasis, we created a *Citrobacter*-induced infectious colitis model. When challenged with *Citrobacter* (**Fig 2E**), GIV-KO mice showed higher fecal bacterial load acutely (**Fig 2F**; 1^st^ wk, 3-fold higher than littermate WT controls), and an unusual delay in bacterial clearance and the development of chronicity (**Fig 2F**; 7^th^ wk, 9-fold higher than littermate WT controls). The GIV-KO mice also displayed colon shortening (**Fig 2G**), patchy transmural chronic inflammation of the small intestine, colon, and rectum (**Fig 2H-I**), and focal muscle hypertrophy and collagen deposits (**Fig 2J-K**). These findings indicate that GIV in myeloid cells is required for effective bacterial clearance in the setting of infection.

### Protective MDP/NOD2 signaling is abolished in myeloid specific GIV-KO mice

Prior studies have shown that pre-treatment with MDP ameliorates infection/bacteremia (<u>Cobb et al. 1986</u>; <u>Matsumoto et al. 1981</u>), fatality in sepsis (<u>Wardowska et al. 2009</u>) and chemical (TNBS and DSS)-induced colitis (<u>Watanabe et al. 2008</u>). We asked if the protective actions of MDP require GIV. Compared to WT controls, the GIV-KO mice developed significantly worse DSS-induced acute colitis (**Fig 3A**), as determined by histological composite scores accounting for deformation of colon crypts and increased immune infiltration in the colon (**Fig 3B-C**) and disease activity index (**Fig 3D-E**); the latter is a composite score of stool consistency, weight loss and the presence of fecal blood (Katkar et al. 2021; Swanson et al. 2020; Whittem <u>et al. 2010</u>). Pre-treatment with MDP ameliorated the severity of colitis in WT, but not GIV-KO mice (**Fig 3B-E**).

**Figure 3.**
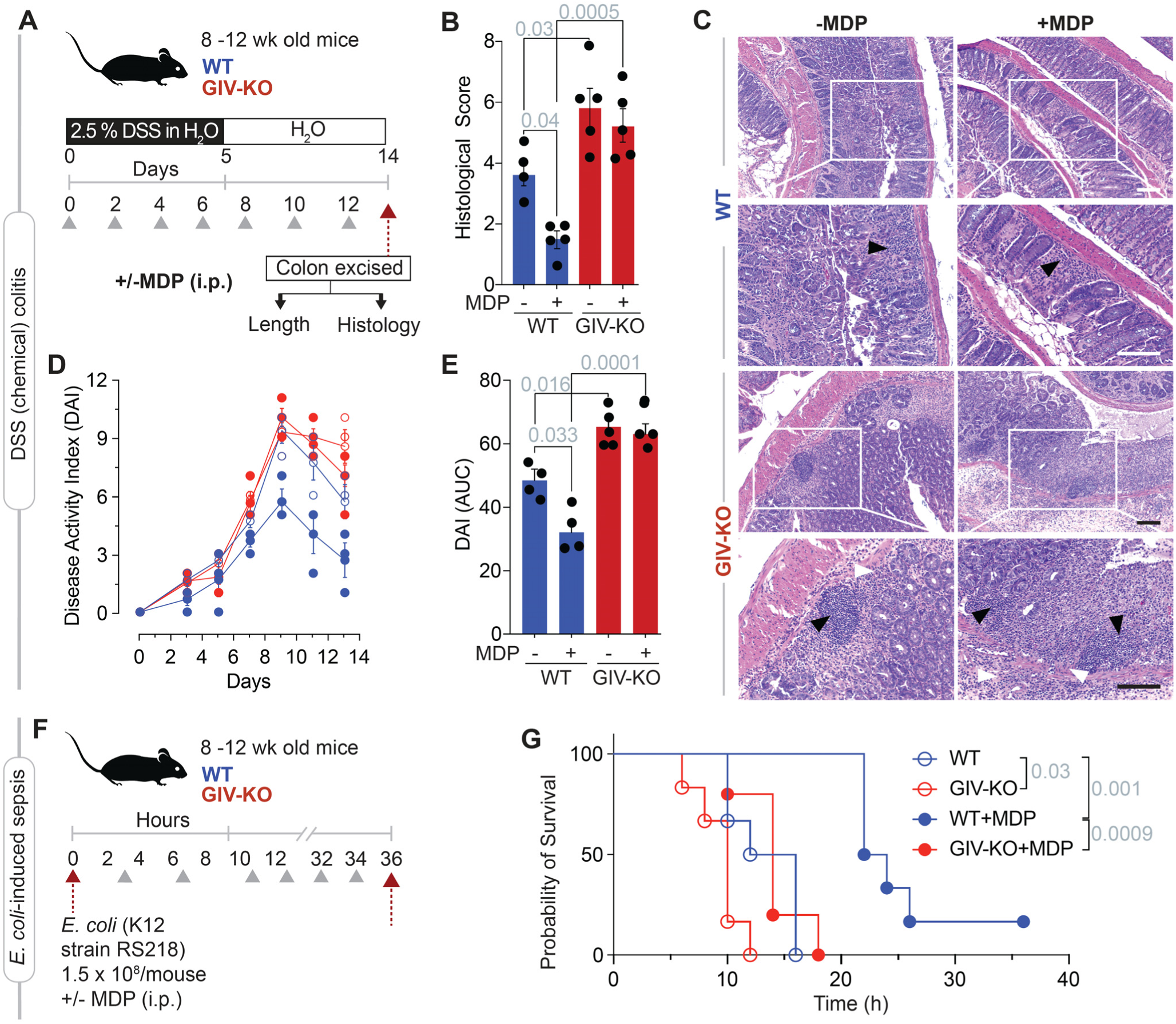
GIV-KO mice are insensitive to the protective actions of MDP/NOD2 signaling. **A-C**. Schematic (**A**) displays the study design for DSS-induced colitis. GIV-KO, n=5; WT, n=5. Findings are representative of two independent repeats. Grey arrowheads denote the alternate day administration of muramyl dipeptide (MDP) (100µg/mouse/day). Bar graphs (**B**) display the histological score, as assessed by a well-accepted methodology (<u>Kim et al. 2012</u>) of analyzing H&E-stained distal colons from the mice. Representative images are displayed in **C**. Arrowheads point to regions of crypt destruction and/or inflammatory infiltrates. Scale bar = 200 µm. **D-E**. Line graphs (**D**) display disease activity index (DAI), calculated for the days 3, 5, 7, 9, 11 and 13 after DSS administration, which accounts for stool consistency (0-4), rectal bleeding (0-4), and weight loss (0-4). Bar graphs (**E**) represent the data in D as area under the curve (AUC). **F-G**. Schematic (F) displays the sepsis study design in which 8 mice in each group were treated with E coli and MDP simultaneously, followed by periodic checks for death (arrowheads). Kaplan-Meier plot (G) displays the % of cohort that survived at those time points. GIV-KO, n=8; WT, n=8. Findings are representative of two independent repeats. *p* values were determined by Mantel-Cox log rank test. *p*-value ≤ 0.05 is considered as significant. All results are displayed as mean ± SEM.

Similar results were seen also in the case of *E. coli*-induced sepsis (**Fig 3F**); fatality was higher in GIV-KO mice compared to WT controls (**Fig 3G**) and pre-treatment with MDP reduced fatality in WT, but not GIV-KO mice (**Fig 3G**). These findings demonstrate that GIV is required for the protective MDP/NOD2 signaling in the setting of infection/inflammation.

### GIV is required for MDP/NOD2-mediated bacterial clearance and controlled inflammation

We next analyzed how depletion of GIV in macrophages impacts MDP/NOD2 signaling. GIV-depleted (shGIV) RAW 264.7 macrophages [a previously-validated model displaying ∼85-90% depletion (<u>Swanson et</u> <u>al. 2020</u>) (**Fig 4A**)] showed significantly higher MDP/NOD2-induced NFkB activity mediated by MDP/NOD2 signaling (**Fig 4B**), as determined using luciferase reporter assays. Findings were confirmed also in HeLa cells (**Fig 4C**), which have used by others for studying NOD2-dependent cellular processes in settings where plasmid transfections are necessary (Damgaard et al. 2012; Hao et al. 2015). HeLa cells depleted or not of GIV (by CRISPR; **Fig S2A**) confirmed that the presence of GIV dampens MDP/NOD2-induced NFkB activity (**Fig 4C**). Thioglycolate-induced primary peritoneal macrophages (TG-PMs) from GIV-KO mice (or their WT littermate controls) stimulated with MDP, proinflammatory cytokines (as detected by ELISA: IL1β, IL6; **Fig 4D** and TNFα, **Fig S2B**) were induced upon MDP stimulation in both WT and KO TG-PMs, but the extent of induction was ∼3-4-fold higher in GIV-KO PMs. Hyperinduction of proinflammatory cytokines was accompanied by a concomitant suppression of the anti-inflammatory IL10 cytokine (**Fig S2B**). These cytokine profiles determined by ELISA generally agreed with gene expression patterns assessed by qPCR (**Fig S2C**). When we infected the TG-PMs with the pathogenic adherent invasive *Escherichia coli* strain-*LF82* (AIEC*-LF82*), which was isolated from CD patients (<u>Darfeuille-Michaud et al. 2004</u>), the KO PMs showed delayed clearance compared to WT PMs (**Fig 4E**). Treatment of the cells with MDP could accelerate clearance in WT, but not KO PMs (**Fig 4E**).

**Figure 4.**
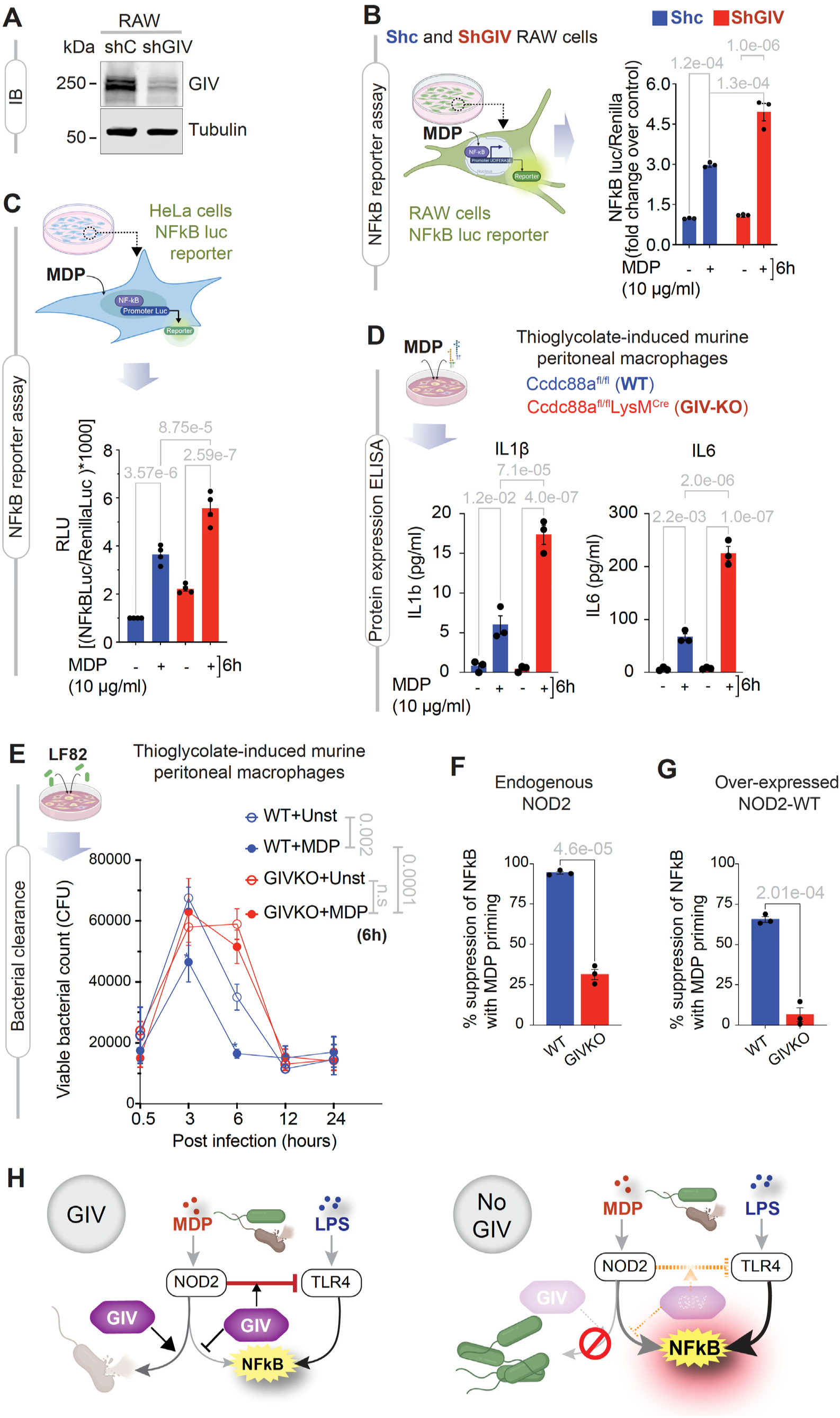
GIV is required for MDP-stimulated bacterial clearance and resolution of inflammation. **A**. Immunoblot of control (Shc) or GIV-depleted (ShGIV) RAW 264.7 cells. **B-C**. Schematics display the NFkB reporter assay in RAW 264.7 (B) and HeLa (C) cells. Bar graphs display the fold change in NFkB activity. **D.** Schematic (top) displays the experimental set up in primary peritoneal macrophages. Bar graphs (bottom) display the concentrations of the indicated cytokines, as determined by ELISA. **E.** Schematic (top) displays the experimental set for bacterial clearance. Line graphs (below) show the viable bacterial counts in the macrophages. **F-G**. Bar graphs display percent suppression of LPS-induced NFkB activity with MDP priming. **H**. Schematic summarizing findings in cells with (left) or without (right) GIV. Statistics: All results are displayed as mean ± SEM (n=3 biological replicates). Significance was tested using two-way/one-way ANOVA followed by Tukey’s test for multiple comparisons. *p*-value ≤ 0.05 is considered as significant.

Prior work has demonstrated that priming of NOD2 with MDP protects cells from excessive inflammation that is induced by LPS (Kim et al. 2015; Watanabe et al. 2008). To test if such protective effect of MDP/NOD2 requires GIV, we used GIVKO and WT HeLa cells. HeLa cells have been used also to study LPS-induced NFkB activation (Hao et al. 2015; Jiang et al. 2017; Manna and Aggarwal 1999; Sulistyowati et al. 2018; Wang et al. 2014; Xiong et al. 2019) and they express MD2, a co-receptor believed to be required for LPS/TLR4 signaling, albeit at low levels (<u>Pridmore et al. 2003</u>). We observed that MDP-pretreatment indeed significantly reduced the degree of NFkB activation in HeLa cells, both in the setting of endogenous NOD2 (**Fig 4F**) and when NOD2 was exogenously overexpressed (**Fig 4G**). In the case of GIV-KO cells, this protective role of MDP was either significantly compromised (**Fig 4F**) or virtually abolished (**Fig 4G**).

Together these findings demonstrate that GIV is required for protective MDP/NOD2 signaling; in its absence, proinflammatory NFkB signaling is excessive (**Fig 4H**). The impact of such protective signaling is more obvious in the MDP-priming studies, in which, the MDP/NOD2→GIV pathway acts antagonistically to the LPS/TLR4 pathway to prevent excessive proinflammatory NFkB signaling (**Fig 4H**).

### GIV is required for the NOD2-dependent induction of key anti-bacterial pathways

NOD2 is known to intricately coordinate numerous intracellular organellar and cytoskeletal processes that facilitate phagocytosis and bacterial clearance (<u>Philpott et al. 2014</u>), e.g., Rac1 activation, actin cytoskeletal rearrangement, autophagy, RIPK activity and the formation of the RIPK2/XIAP aggrosomes (‘RIPosome’ (Ellwanger et al. 2019; Gong et al. 2018)). To predict which of these processes are impacted by GIV, we resorted to protein-protein interaction (PPI) network analysis. First, a PPI network was built using previously published NOD1, NOD2 and GIV interactomes, as determined using proximity-based biotinylation assay, BioID (see *Methods*; **Fig 5A**). Then we carried out *in silico* perturbation to either delete GIV (to mimic GIV-KO/shGIV cells) or selectively remove the edge connecting NOD2 and GIV (to mimic binding-deficient NOD2/GIV mutants) (**Fig 5A; S3A-C**). Topological network-based differential analysis of perturbed vs unperturbed networks (see *Methods*) showed that NOD2’s connectivity with many proteins (‘nodes’) and pathways were altered in the perturbed state (detailed in **Fig S4**, **S3A-C**; summarized in **Fig 5B**) and would result in a broad derailment of the functions of NOD2, ranging from cytoskeletal remodeling, through NFkB and RIPK2 signaling, to ATG16L1-dependent autophagy.

**Figure 5:**
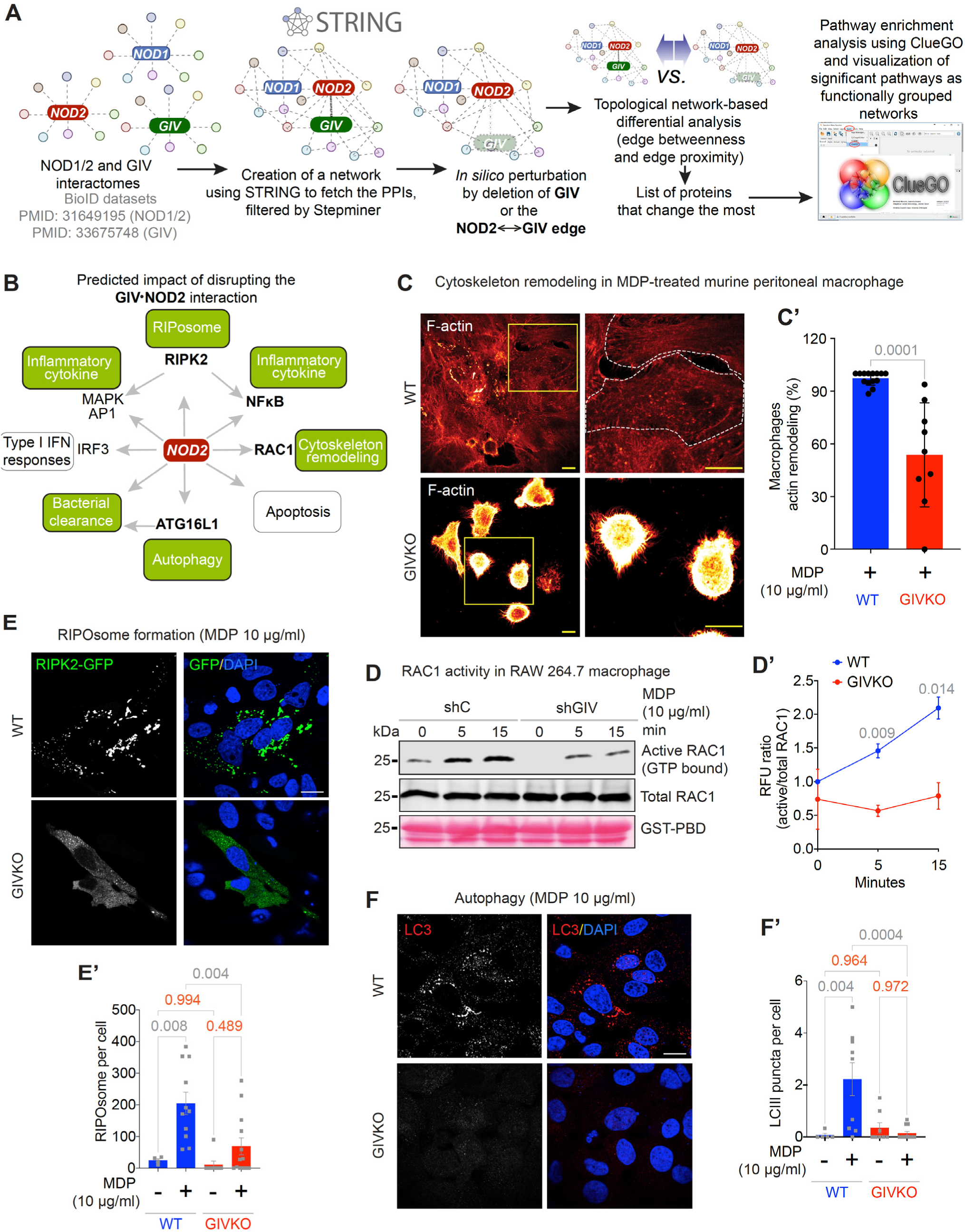
GIV is required for NOD2-dependent anti-bacterial cellular processes. **A.** Schematic summarizes the key steps in building a protein-protein interaction (PPI) network using NOD1, NOD2 and GIV interactomes as seeds and connecting them with the shortest path prior to network analysis. *In silico* perturbation of the network (see **Supplementary** Figure 3A) was performed by either deleting GIV (see **Supplementary** Figure 3B) or selectively removing the interaction between GIV and NOD2 (see **Supplementary** Figure 3C). The original and perturbed networks were then analyzed for topological network-based differential analyses to identify the impact of these perturbations on the connectivity of NOD2 with other proteins in the network. The list of impacted proteins was analyzed by ClueGo to identify the cellular pathways and processes that were significantly enriched, and hence, impacted by the perturbations. **B.** Schematic summarizing the known major NOD2-associated cellular processes; highlighted in green are those that were predicted to be impacted significantly. Major hub proteins that regulate these processes and are predicted to be impacted by PPI network perturbation. See **Supplementary** Figure 4A-C for detailed PPI network analyses. **C-C’**. Thioglycolate-stimulated murine peritoneal macrophages (TGPMs) isolated from both WT and GIVKO mice were stimulated with MDP (10 µg/ml) for 1h at 37°C and stained with phalloidin, a F-actin stain. Representative images (C) are shown. See also **Supplementary** Figure 5A for the untreated controls for WT and GIV-KO cells. Quantification of images for % macrophage with defective actin organization is displayed as scattered plot with bar graph (C’; ∼30-40 cells/assay; n = 3 mice in each group). Scale bar = 10 µm. **D-D’**. Immunoblot (D) of GIV-depleted or control RAW 264.7 macrophages stimulated with MDP (10 µg/ml) for 0, 5 and 15 min, and probed for Rac1 activation by pulldown assays using GST-PBD. Line graph (D’) displaying RFU quantification of the same. **E-E’**. GIV depleted or control HeLa cells expressing GFP-RIPK2 were stimulated, or not, with MDP (10 µg/ml) for 6 h at 37°C prior to fixation and staining (green = GFP; blue, DAPI/nuclei) and analyzed by confocal microscopy. Representative images (E) are shown. Scale bar = 10 µm. See also **Supplementary** Figure 5B for black and white images of the green channel and the untreated controls for WT and GIV-KO cells. Quantification of images using particle count application (ImageJ), representing RIPosome counts per cell is displayed as bar graph (E’). (∼30-40 cells/assay; n = 3 mice in each group). **F-F’.** GIV depleted or control HeLa cells were treated as in 5E before staining for the marker of autophagosome, LC3 (red) and DAPI (blue; nuclei) and analyzed by confocal microscopy. Representative images (F) are shown. Scale bar = 10 µm. Quantification of images, carried out as in 5E, representing autophagosomal structures per cell is displayed as bar graph (F’). (∼30-40 cells/assay; n = 3 mice in each group). See also **Supplementary** Figure 5C for the untreated controls for WT and GIV-KO cells. All results are displayed as mean ± SEM. Statistical significance was determined using unpaired t-test. *p*-value ≤ 0.05 is considered as significant.

Having observed earlier that GIV depletion induced a hyperinflammatory response and impaired bacterial clearance (**Fig 4**), we sought to validate some of the other predictions. MDP-stimulated actin cytoskeletal remodeling (determined by Phalloidin staining; **Fig 5C-C’**; **Fig S5A**) and Rac1 activity (determined by GST-PBD pulldown assays; **Fig 5D-D’**) were significantly impaired in GIV-depleted TG-PMs. In the absence of reliable phosphoantibodies to assess RIPK2 activity, we resorted to a published and well-characterized microscopy-based quantifiable reporter assay; it involves monitoring ligand-dependent EGFP-RIPK2 redistribution in HeLa cells from diffuse cytosolic to bright punctate macromolecular aggregates (<u>Ellwanger et al. 2019</u>), a.k.a RIPosomes. Such redistribution requires NOD2 activation and follows NFkB activation (but does not depend on it). MDP stimulated the assembly of RIPosomes in WT cells, but significantly reduced in GIV-KO cells (**Fig 5E-E’**; **Fig S5B**); where assembled, GIV colocalized with the RIPosomes (**Fig S5D-E**). These findings indicate that GIV, a *bona fide* actin remodeler (<u>Enomoto et al. 2006</u>), is required for the assembly of RIPosomes, which is consistent with prior work showing that RIPosome assembly shifts EGFP-RIPK2 from triton-soluble to insoluble fractions and that these structures co-localize with actin but not membrane compartments (endosomes/autophagosomes) (<u>Ellwanger et al. 2019</u>).

Because an intact NOD2→RIPK2 signaling axis initiates autophagy in an ATG16L1-dependent manner (<u>Cooney et al. 2010</u>), and NOD2 connectivity to both RIPK2 and ATG16L1 is predicted to be impacted in the perturbed networks (**Fig 5B**; **Fig S3C-D**), we assessed autophagy in the same cells as above. MDP-stimulated autophagy was induced in WT, but not GIV-KO cells, as determined by a quantitative assessment of the abundance of punctate LC3-positive vesicles/cell (**Fig 5F-F’**). These findings indicate that GIV is required for the induction of the NOD2→RIPK2→ATG16L1 cascade that culminates in the initiation of autophagic flux. Because this cascade is an autophagy-dependent, anti-bacterial pathway implicated in CD pathogenesis (<u>Homer et al. 2010</u>), its failed induction is consistent with impaired bacterial clearance in GIV-KO cells we observed in **Fig 4E**.

### The GIV●NOD2 interaction is direct and is dynamically regulated by MDP, ATP, pH and receptor dimerization

NOD2 exists in an inactive, ADP-packed form through stabilized intramolecular interactions; upon MDP binding, conformational changes allow ADP→ATP exchange, self-oligomerization, and downstream signaling (Ellwanger et al. 2019; Gong et al. 2018) (**Fig 6A**). We asked if/how these sequential events may regulate GIV’s ability to bind NOD2. Using co-immunoprecipitation studies on lysates expressing both proteins as full length, we first validated the BioID findings (<u>Lu et al. 2019</u>), i.e., GIV preferentially binds NOD2, but not NOD1 (**Fig 6B**). Next we carried out co-immunoprecipitation assays at various time points after MDP stimulation, using either HA-NOD2 (**Fig 6C**) or GIV-FLAG (**Fig 6D**) as bait, and assess GIV-FLAG and HA-NOD2, respectively, as preys. Regardless of the bait-prey pairing, co-complex assembly was induced in both instances within ∼1 h, sustained up to ∼3 h, followed by disassembly by ∼6 h after MDP stimulation (**Fig 6C-D**). To compare MDP-induced co-complexes against those observed with specific nucleotide-bound forms of NOD2, we treated permeabilized cells with ADP, ATP or its non-hydrolyzable ATPγS analog prior to their use as source of protein in co-immunoprecipitation assays (**Fig 6E**-*left*). ATP and ATPγS-loaded NOD2 showed a similar degree of co-complex assembly with GIV as observed with MDP, whereas, binding was abrogated in the setting of ADP (**Fig 6E**-*right*). Findings indicate that the NOD2●GIV interaction is augmented by MDP and the ATP-bound active conformation, two intertwined events during receptor activation (<u>Maekawa et al. 2016</u>).

**Figure 6.**
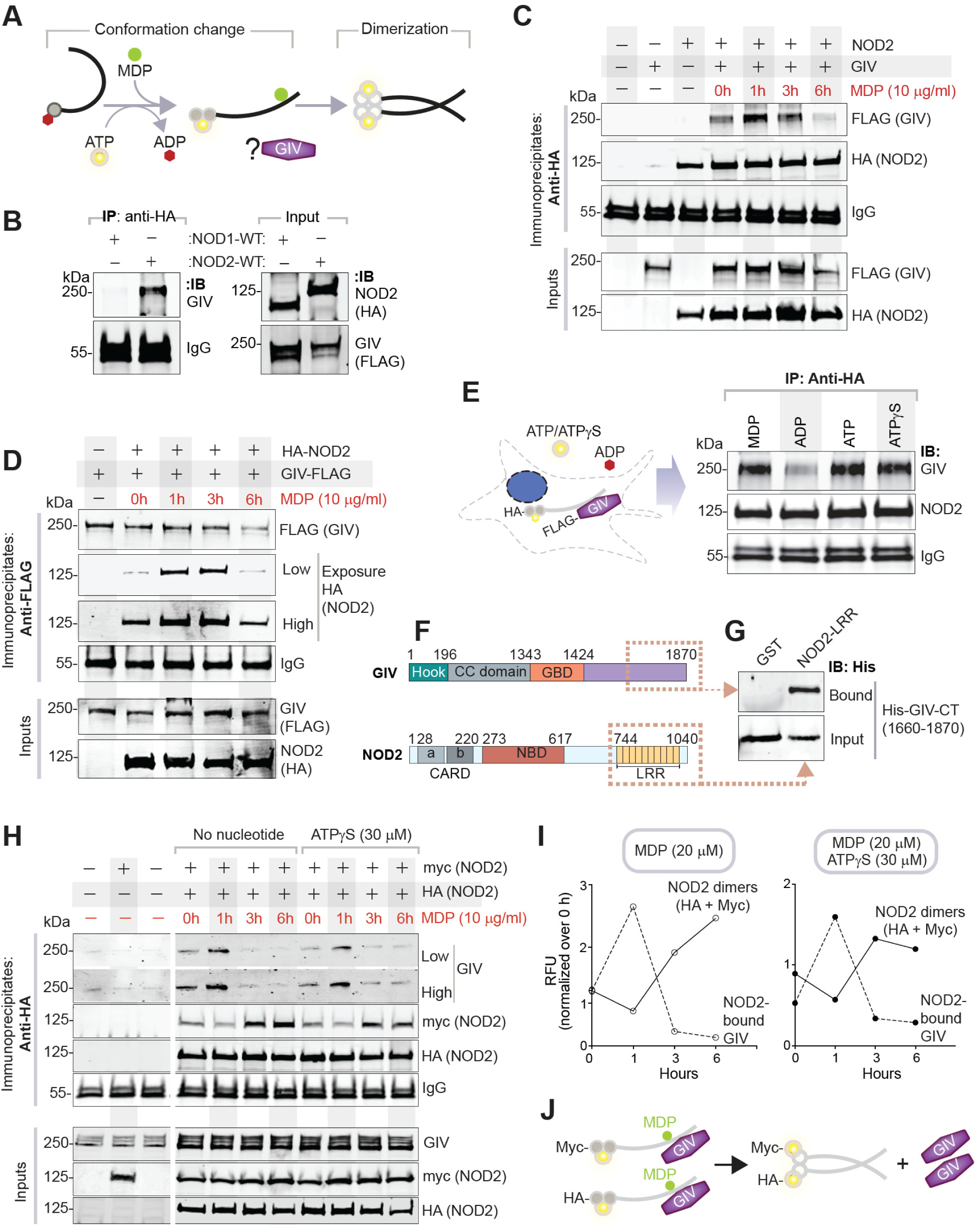
The NOD2(LRR)●GIV(C-term) interaction is direct and dynamically regulated. **A.** Schematic displays key steps in MDP-induced NOD2 signaling. In resting cells, ADP-bound inactive NOD2 exists in an autoinhibited conformation. Upon ligand (MDP) stimulation, ADP is exchanged for ATP, which further enhances MDP-binding and is associated with conformational change and ‘opening’ of the LRR module. Subsequently, NOD2 dimerizes and forms complexes with other signaling proteins. **B.** HA-tagged NOD1/2 proteins were immunoprecipitated from equal aliquots of lysates of HEK cells co-transfected with GIV and either NOD1 or NOD2 using anti-HA mAb. Immunoprecipitated (IP; left) complexes and input (right) lysates were analyzed for NOD1/2 and GIV by immunoblotting (IB). **C.** HA-tagged NOD2 was immunoprecipitated with anti-HA mAb from equal aliquots of lysates of HEK cells expressing GIV-FLAG and HA-NOD2, stimulated (+) or not (-) with MDP for indicated time points. Immunoprecipitated (IP; top) complexes and input (bottom) lysates were analyzed for NOD2 and GIV by immunoblotting (IB). **D.** FLAG-tagged GIV was immunoprecipitated with anti-FLAG mAb from equal aliquots of lysates of HEK cells expressing GIV-FLAG and HA-NOD2, stimulated (+) or not (-) with MDP for indicated time points. Immunoprecipitated (IP; top) complexes and input (bottom) lysates were analyzed for NOD2 and GIV by immunoblotting (IB). **E.** HA-tagged NOD2 was immunoprecipitated with anti-HA mAb from equal aliquots of lysates of HEK cells as in **6C**, either after stimulation with MDP (1 h; lane 1) or after permeabilization and treatment with ADP, ATP or ATPγS for 30 min. Immune complexes were analyzed for GIV and NOD2 by IB. **F-G**. Schematic (F) depicts the domain maps of NOD2 and GIV. Boxed regions indicate the domains that were used to generate His- and GST-tagged recombinant proteins for use in GST-pulldown assays in G. Equal aliquots of recombinant His-GIV-CT (∼3 µg; input, bottom) was used in pulldown assays with immobilized GST and GST-NOD2-LRR. Bound (top) GIV was visualized by immunoblot (IB). **H-J**. HA-tagged NOD2 was immunoprecipitated with anti-HA mAb from equal aliquots of lysates of HEK cells co-expressing HA and myc-tagged NOD2 and GIV-FLAG, permeabilized and exposed (or not) to ATPγS for 30 min prior to stimulation (+) or not (-) with MDP for indicated time points at 37°C. Immunoprecipitated (IP; H, top) complexes and input (H, bottom) lysates were analyzed for HA/myc-tagged NOD2 and GIV by immunoblotting (IB). Line graphs (I) display the quantification of representative immunoblots using band densitometry by LiCOR Odyssey. Schematic (J) summarizes the inference drawn from the IP studies; disassembly of NOD2•GIV complexes coincides with the assembly of NOD2 dimers.

Because GIV’s ∼210 aa C-term (CT) is the module that consistently couples to diverse ‘receptors/sensors (<u>Ghosh and Mullick 2021</u>)’ via short linear motifs (SLIMs), we asked if GIV-CT is sufficient to bind NOD2. Recombinant GST-GIV-CT bound full length NOD2 but not NOD1 from lysates (**Fig S6A**). Using recombinant proteins, we confirmed that the C-term of GIV directly binds NOD2-LRR (**Fig 6F-G**, **Fig S6B**). Because MDP binds NOD2-LRR with high affinity at a pH range from 5.5 to 6.5 (<u>Grimes et al. 2012</u>), we tested if pH and MDP impact GIV’s ability to bind NOD2-LRR. Binding of GIV was progressively increased at lower pH range of 5.5 to 6.5 (<u>Grimes et al. 2012</u>) (**Fig S6C**), and remained unchanged regardless of the presence or absence of molar excess of MDP (**Fig S6C-D**). Furthermore, we assessed how the dynamics of assembly of NOD2●GIV complexes coincide with the assembly of NOD2 oligomers, by measuring the relative abundance of MDP-stimulated GIV•NOD2 and differentially-tagged NOD2(myc)•NOD2(HA) co-complexes in immunoprecipitation studies. The disassembly of NOD2●GIV complexes coincided with the assembly of NOD2•NOD2 di/oligomers (**Fig 6H-I**).

These findings demonstrate that the C-term of GIV directly binds the LRR domain of NOD2. This interaction is both dynamic and finite (see **Fig 6J**); assembly is initiated by ligand stimulation and ATP-bound active conformation and is favored by low pH, and that disassembly coincides with NOD2 di/oligomerization.

### GIV binds the terminal LRR of NOD2, cannot bind the *1007fs* variant which lacks the terminal LRR

We next asked if/how binding of GIV maybe impacted by 3 NOD2 variants (*R702W*, *G908R* and *1007fs*) that represent ∼80% of the mutations independently associated with susceptibility to CD (<u>Lesage et al. 2002</u>). The affected aa are located either proximal to or within the LRR domain (**Fig 7A**). Binding to GIV was impaired in the case of the two mutations (*R702W* and *1007fs*; **Fig 7B**) which demonstrate the highest disease penetrance (∼100%; **Supplementary Table 1**). The MDP-contact site variant (<u>Boyle et al. 2013</u>; <u>Vijayrajratnam et al. 2017</u>) (*G908R*) bound GIV to similar extent as NOD2-WT (**Fig 7B**). Further studies with the *1007fs* variant confirmed that it remains binding-deficient even upon MDP stimulation (**Fig 7C**). Co-immunoprecipitation (**Fig S6E**) and GST-pulldown studies with GIV-CT (**Fig 7D**, **Fig S6F**) were in agreement when both approaches revealed that binding is reduced upon mutagenesis of key Arg residues (*R1034* and *R1037*) that stabilize the terminal LRR. These studies also showed that binding is virtually abolished upon deletion of the terminal repeat in the LRR domain (Δdel) (**Fig 7D**; **Fig S6E-F**). GST pulldown assays with GST-LRR and His-GIVCT recombinant proteins further confirmed that the NOD2-LRR Δdel mutant (which lacks the 10^th^ LRR repeat exactly like the naturally occurring *1007fs* variant; **Fig 7A**) loses its ability to bind GIV (**Fig 7E**). Findings demonstrate the requirement of the terminal LRR repeat in NOD2 for binding GIV (**Fig 7F**).

**Figure 7:**
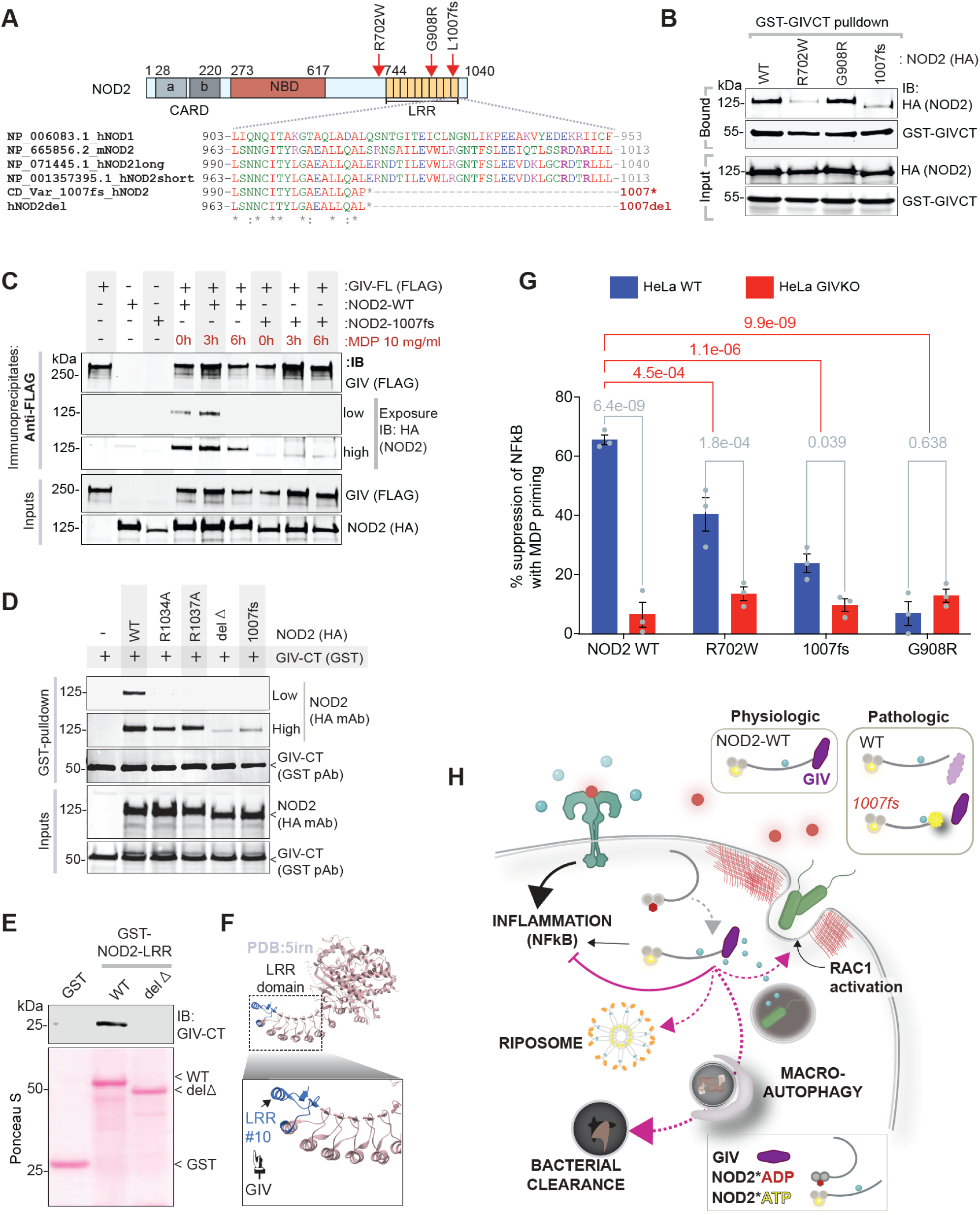
Characterization of the NOD2•GIV interface exploiting CD associated NOD2 mutants. **A.** *Top*: Bar diagram showing domains of NOD2. *Bottom*: An alignment of the amino acid sequence of the 10^th^ LRR repeat of human (h) and mouse (m) NOD2 with hNOD1 and the CD-risk associated NOD2 variant (NOD2-1007fs) and the deletion mutant generated in this work (NOD2-Del) is shown. Residues mutated in this study to evaluate potential participating residues in the NOD2•GIV interaction are highlighted. **B.** GST-GIV-CT was pulled down using Glutathione beads from equal aliquots of lysates of HEK lysates co-expressing GST-GIV-CT (aa 1660-1870; mammalian p-CEFL vector) and wild-type (WT) or 3 indicated CD-risk associated variants of HA-NOD2. Bound NOD2 proteins and similar expression of GIV-CT was assessed by immunoblotting (IB) using anti-HA (NOD2) and anti-GST (GIV-CT) antibodies. **C.** FLAG-tagged GIV was immunoprecipitated with anti-FLAG mAb from equal aliquots of lysates of HEK cells expressing GIV-FLAG and either wild-type (WT) or *1007fs* variant of HA-NOD2, stimulated (+) or not (-) with MDP for indicated time points. Immunoprecipitated (IP; top) complexes and input (bottom) lysates were analyzed for NOD2 and GIV by immunoblotting (IB). **D.** GST-GIV-CT was pulled down using Glutathione beads from equal aliquots of lysates of HEK lysates co-expressing GST-GIV-CT (aa 1660-1870; mammalian p-CEFL vector) and either WT or HA-NOD2 mutants predicted to disrupt NOD•GIV binding. Immunoprecipitated (IP; top) complexes and input (bottom) lysates were analyzed for NOD2 and GIV-CT by immunoblotting (IB) as in **7B**. **E-F**. GST-pulldown assays (E) carried out as in B, using GST and NOD2-LRR proteins as indicated and bound His-GIVCT is assessed. Schematic (F) highlights the terminal LRR repeat (blue) of NOD2 which binds GIV. **G.** Bar graphs display NFkB reporter assay in HeLa cells. Cells were pre-incubated with MDP (1µg/ml) and then stimulated with LPS (100ng/ml) and % suppression of NFkB activity was detected using dual cell reporter assay. **H.** *Left*: Schematic summarizing key findings in this work. Magenta colored solid and interrupted lines indicate the GIV-dependent impact on NOD2 that were interrogated in this work. *Right*: In physiology, bacterial sensing and signaling by NOD2 requires GIV to limit inflammation. In pathology, dysregulated inflammation results when either wild-type NOD2 cannot bind GIV (e.g., GIV is low/absent) or when CD-risk associated *1007fs* variant cannot bind GIV.

We next analyzed how the NOD2 CD-risk variants respond to LPS when pre-treated with MDP. To specifically assess the role of GIV, we assessed each mutant in NFkB luc assays in cells with or without GIV. In the presence of NOD2-WT, MDP-pretreatment significantly suppressed the degree of LPS-stimulated NFkB activation; however, that degree of protection was impaired in the case of the NOD2 mutants (**Fig 7G**; red *p*-values). For example, compared to NOD2-WT, the NOD2-*1007fs* variant that cannot bind GIV shows an impairment in protective signaling from ∼65% to ∼20% (**Fig 7G**). In the case of GIV-KO cells, the degree of suppression by NOD2-WT was reduced from ∼65% to ∼10% (**Fig 7G**; gray *p*-values). The fact that ∼10-20% reduction occurs in settings where GIV cannot bind NOD2 (10% in GIV-KO or 20% in NOD2-*1007fs*) suggest that there could be potential GIV-independent mechanisms or endogenous NOD2 in the cells. Regardless, the findings indicate that the GIV●NOD2 interaction contributes to a significant part of the protective role of MDP.

## DISCUSSION

The major discovery we report here is the identification of a direct binding partner of NOD2 that has a dual role upon sensing bacteria and/or bacterial products (**Fig 7H**): (i) it suppresses NFkB-dependent inflammatory signals, and (ii) concomitantly enhances anti-bacterial pathways and processes that are known to be NFkB independent. Such dual function protects the host against exuberant inflammation in the setting of infections. We define the molecular basis for the actions of GIV and demonstrate the consequences when those actions are is perturbed. Insights gained spur new paradigms in innate sensing and signaling via NOD2 in three ways.

### GIV enhances tolerant (protective) NOD2 signaling in macrophages

NOD2-mediated dampening of NFkB serves as an important, but poorly understood protective mechanism (<u>Ashton et al. 2022</u>). NOD2 also protects host cells by facilitating bacterial clearance by coordinating numerous cytoskeletal, organellar and signaling processes and pathways. By demonstrating that binding of GIV to NOD2 is critical for both arms of the protective response, we provide hitherto unforeseen insights into how NOD2 coordinates macrophage responses via measured activation of inflammation that is optimal for immunity and tissue homeostasis. In the absence of GIV, the protective functions of NOD2 are lost, microbes are not cleared efficiently and macrophages assume a reactive phenotype. These findings are in keeping with our previous work, in which we demonstrated how GIV serves as a ‘brake’ within the LPS/TLR4 siganling cascade (<u>Swanson et al. 2020</u>). Using a conserved motif in its C-terminus, GIV also binds and prevents inflammatory signaling of other TLR receptors including TLR1/2, TLR2/6 and TLR3 (<u>Swanson et al. 2020</u>). Therefore, GIV’s role in two major classes of pattern recognition receptors (PPRs), NOD2 and TLRs, is to induce tolerogenic programs that are geared towards tissue homeostasis and immunity. In doing so, GIV emerges as a shared point of convergence for major PPRs that coordinate early host defense against pathogen invasion (Lu et al. 2019; Swanson et al. 2020; Zhou et al. 2018). Whether GIV serves as a platform for crosstalk between the two PRRs requires further dissection.

### The GIV●NOD2 interaction couples new pathways to NOD2

We show that the GIV●NOD2 interaction is specific (no binding to NOD1), direct, and is dynamically regulated by a multitude of factors that are intricately associated with receptor activity (favored by MDP and ATP), conformation (favored by ATP, but abolishd in oligomeric state). That assembly is favored at pH 5.5-6.5 suggests that, like described earlier for MDP (<u>Grimes et al. 2012</u>), the *in vivo* interaction between GIV and NOD2-LRR may happen in acidic compartments. Binding is mediated by GIV’s C-terminal ∼210 aa which interacts with the terminal LRR repeat of NOD2; this makes GIV the 3^rd^ such interactor of NOD2-LRR (only 2 prior) (<u>Boyle et al. 2014</u>), and the only one that suppresses NFkB to augment protective signaling. Although we reveal here how GIV impacts NOD2 signaling, if/how NOD2 may impact GIV’s functions as a cAMP inhibitor (Getz et al. 2019; Ghosh et al. 2017) (via Gi activation and Gs inhibition) remains unknown. Because high cAMP favors NOD2-mediated clearance of microbes (<u>Kim et al. 2016</u>), the GIV●NOD2 interaction could serve as means to abrogate GIV-dependent cAMP inhibition. If that is the case, NOD2 will join the list of ligand/receptor pairs that modulate trimeric GTPase proteins and cellular cAMP (<u>Ghosh and Mullick 2021</u>) by accessing the C-term of GIV. In all these instances, each receptor/sensor interacts with SLIMs within the C-terminus of GIV (<u>Ghosh and Mullick 2021</u>); the identity of the NOD2-binding SLIM and if it overlaps with the TLR4- or Gαi/s-binding SLIMs remains unknown. These directions will be explored next.

### Loss of NOD2●GIV interaction is a feature of the 1007fs CD-risk variant

The three main CD-associated variants (*R702W*, *G908R*, and *1007fs*) all contribute to disease pathogenesis through interference with bacterial recognition (<u>Heerasing and Kennedy 2017</u>). While one (*G908R*) has clear structural basis for impaired MDP contact site (Boyle et al. 2013; Vijayrajratnam et al. 2017), the others (*R702W* and *1007fs*) show defects in palmitoylation and impaired PM localization (<u>Lu et al. 2019</u>). However, restoring PM localization rescues the *R702W* and *G908R* variants, but not the *1007fs* variant (<u>Lécine et al.</u> <u>2007</u>), indicating that the *1007fs* variant lacks key functions, perhaps because of the truncated terminal LRR repeat. We show that the *1007fs* variant does not bind GIV and that it lacks the same terminal LRR repeat that is essential for the NOD2●GIV interaction. We confirmed this not just in co-immunoprecipitation studies, but also *in vitro* in pulldown assays using recombinant NOD2-LRR proteins; the pulldown studies rule out the possibility that loss of GIV binding we observe in co-immunoprecipitation studies is merely a consequence of protein mislocalization in cells, as implicated previously for other NOD2-interactors, e.g., Erbin (<u>McDonald et</u> <u>al. 2005</u>). Thus, GIV emerges as a first-in-class NOD2-interactor that requires the 10^th^ LRR repeat, which is lost in the *1007fs* variant. Because this variant (also termed *3020insC*) is most consistently associated with CD across multiple studies and population groups (<u>Economou et al. 2004</u>), and displays 100% disease penetrance, it is not surprising that our GIV-KO animals challenged with *Citrobacter* develop key features of CD, e.g., patchy ileocolitis with transmural inflammation, focal muscle hypertrophy and fibrosis, and dysbiosis with defective bacterial clearance. It is noteworthy that these developed within 7 wks, which is relatively early compared to the only other known murine model of CD SAMP1/YitFcs, which takes ∼30 wks (<u>Pizarro et al.</u> <u>2011</u>). The latter develop CD-like spontaneous ileitis, via genetic defects in many genes, e.g., *Pparγ, Nod2, Il10ra*, among others. Future studies will assess whether *Citrobacter*-challenged GIV-KO mice progress to recapitulate some or most features of CD at a molecular (defective innate and adaptive immune response) and phenotypic (develop fistula) level.

## LIMITATIONS OF STUDY

Although this work established phenotypes in GIV-KO mice and cultured cells, the importance of the GIV●NOD2 interaction in the phenotypes could not be interrogated due to the lack of specific tools; to do so, identification of precise binding-deficient GIV and/or NOD2 mutants is a must. It is noteworthy that whenever possible, all phenotypes were assessed in the context of MDP stimulation, which helps maintain the specificity of the readouts and conclusions drawn.

## Acknowledgments

This work was supported by the National Institutes of Health (NIH) grant R01-AI141630 (to PG). PG and DS were also supported by the Leona M. and Harry B. Helmsley Charitable Trust and the NIH (UG3TR003355, UG3TR002968 and R01-AI55696). G.D.K. was supported through The American Association of Immunologists Intersect Fellowship Program for Computational Scientists and Immunologists. Other sources of support include--T32GM8806 (to D.V.). S-RI was supported by the postdoctoral fellowship grant from NIH (3R01DK107585-02S1). We are grateful to Soumita Das (UCSD) for resources and knowhow for bacterial studies and constructs. We thank Ying Jones (UC San Diego Electron Microscopy Core Facility) for technical and logistical support. We also thank Lee Swanson, Ibrahim Sayed for their comments, feedback, and technical support. This manuscript includes data generated at the UC San Diego Institute of Genomic Medicine (IGM) using an Illumina NovaSeq 6000 that was purchased with funding from a National Institutes of Health SIG grant (#S10 OD026929). The content is solely the responsibility of the authors and does not necessarily represent the official views of the Helmsley Charitable Trust or the National Institutes of Health.

## Author contributions

G.D.K, M.A and P. G. conceptualized the project. G.D.K., M.A., S.S, J.C, Y.M, C.R.E, M.M and V.C, were involved in data curation and formal analysis. D.T.V, with supervision from D.S, carried out 16S microbiome analysis. S.S, with supervision from P.G carried out the protein-protein network analyses. G.D.K conducted all animal studies. M.A conducted all biochemical studies. V.C assisted G.D.K. for ELISA. Y.M assisted G.D.K. for qPCR and image analyses. C.R.E and S-R.I assisted M.A in construct design, cloning and mutagenesis. G.D.K. and P.G. prepared figures for data visualization, wrote original draft, reviewed and edited the manuscript. P.G. supervised and acquired funding to support the study. All co-authors approved the final version of the manuscript.

## Inclusion and diversity

We worked to ensure sex balance in the selection of non-human subjects. One or more of the authors of this paper belongs to a minority group underrepresented in science.

## KEY RESOURCE TABLE

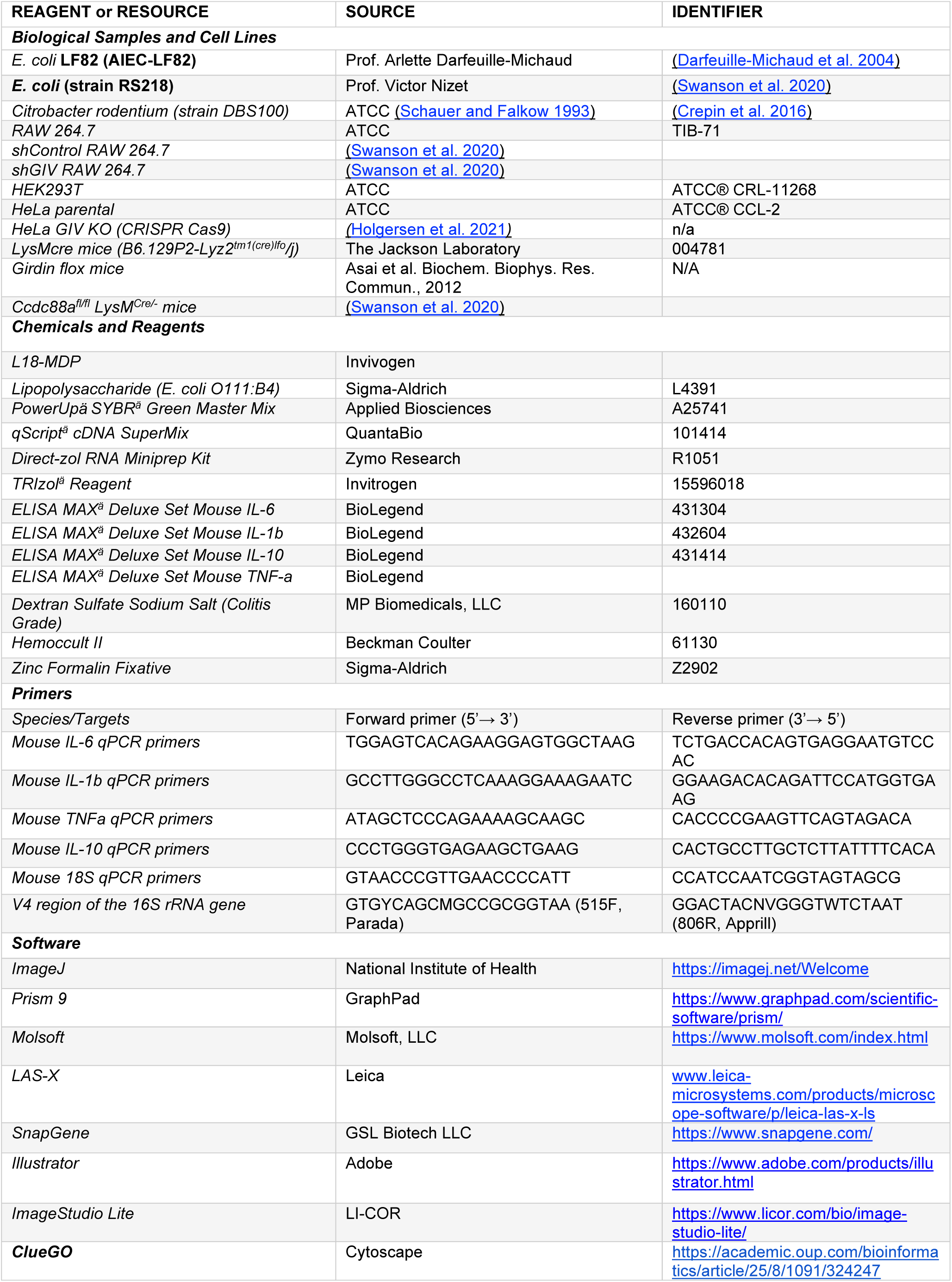
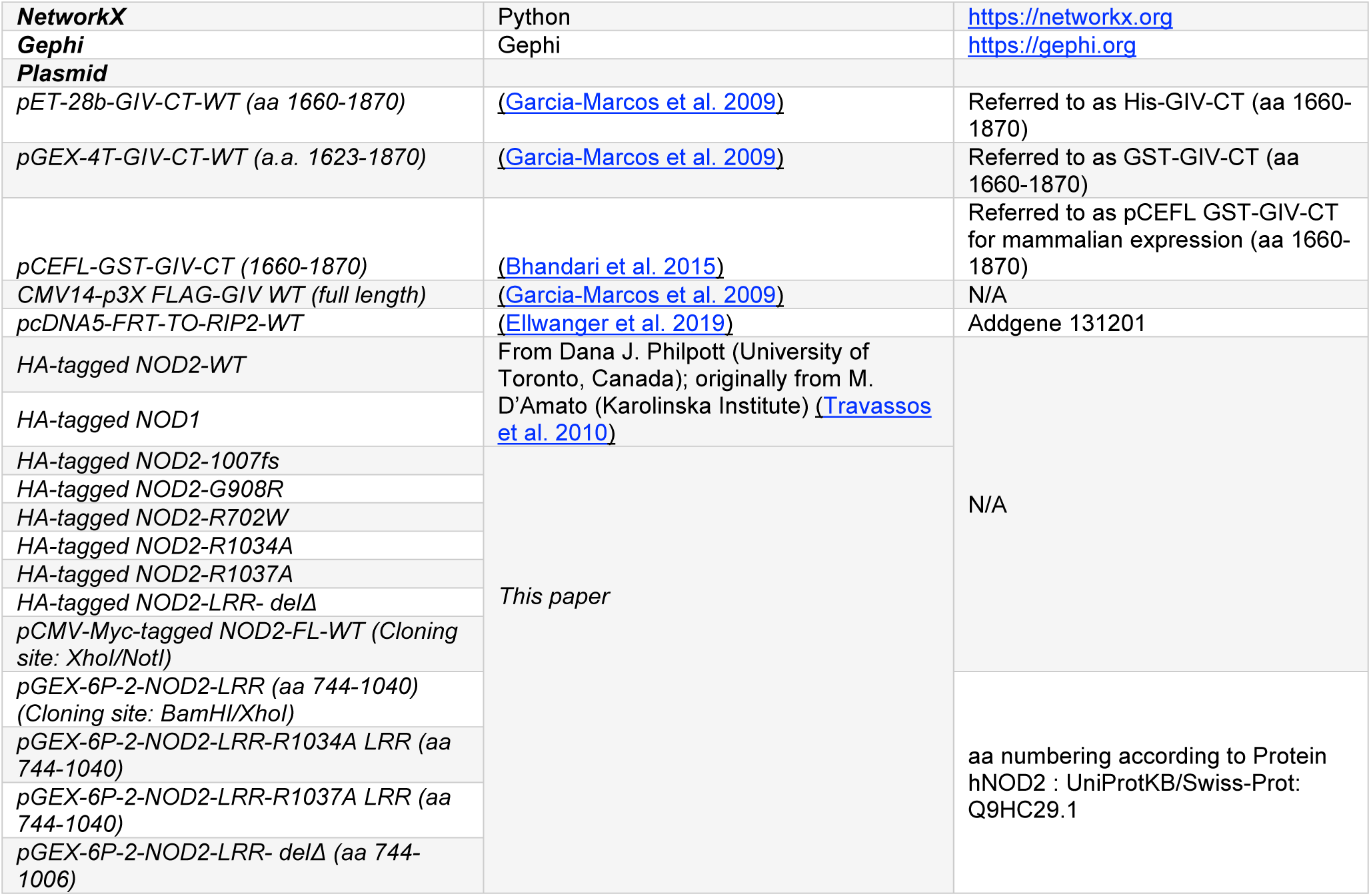

**Contact for Reagent and Resource Sharing** Pradipta Ghosh

## DETAILED METHODS

### Cell culture

Thioglycolate-elicited murine peritoneal macrophages (TGPMs) were collected from peritoneal lavage of 8- to 12-wk-old C57BL/6 mice with ice cold RPMI (10 ml per mouse) 4 days after intraperitoneal injection of 3 ml of aged, sterile 3% thioglycolate broth (BD Difco, USA) and cultured as described previously (<u>Pavlou et</u> <u>al. 2017</u>). Cells were passed through 70 µm filter to remove possible tissue debris contamination during harvesting. Cells were counted, centrifuged, and resuspended in RPMI-1640 containing 10 % FBS and 1% penicillin/streptomycin. Cells were plated with required cell density and the media was changed after 4 h to remove non adherent cells. Cells were allowed to adjust to overnight culture before the addition of stimuli as indicated in the figure legends. RAW264.7, HEK293 and HeLa cells (from ATCC) were maintained in the DMEM media containing 10% FBS and 1% penicillin/streptomycin.

### Bacteria and bacterial culture

*Citrobacter rodentium* (strain DBS100) and Adherent Invasive *Escherichia coli* strain LF82 (AIEC-LF82) were cultured from a single colony inoculation into LB broth for 6-8 h on shaking incubator, followed by overnight culture under oxygen-limiting conditions, without shaking, to maintain their pathogenicity as done previously (Das et al. 2011; Sarkar et al. 2017; Sayed et al. 2020). Bacterial cells were counted by measuring an absorbance at 600 nm (OD600), washed with PBS, and infected with indicated MOI in figure legends.

### Mice

*Ccdc88a^fl/fl^* mice were a gift from Dr. Masahide Takahashi (Nagoya University, Japan) and was developed as described (<u>Asai et al. 2012</u>). LysM*^Cre/Cre^* mice (B6.129P2-Lyz2tm1(cre)lfo/j) were purchased from the Jackson Laboratory. *Ccdc88a^fl/fl^* x LysM*^Cre/-^* mice were generated previously by us as described (<u>Swanson et al. 2020</u>) and were maintained as homozygous floxed and heterozygous LysMcre. Primers required for the genotyping are mentioned in primer table. All mice studies were approved by the University of California, San Diego Institutional Animal Care and Use Committee (IACUC). Both male and female mice (8-12 weeks) were used and maintained in an institutional animal care at the University of California San Diego animal facility on a 12-hour/12-hour light/dark cycle (humidity 30–70% and room temperature controlled between 68–75 °F) with free access to normal chow food and water.

### Fecal pellet collection, DNA extraction and 16S rRNA sequencing

Individual mice fecal pellets from GIV-KO mice and their littermate WT controls (8-12 weeks) were collected in clean containers and frozen at -80°C untill use. Samples were transported to microbiome core facility, University of California, San Diego for 16S rRNA processing. Total DNA was extracted from the individual mice fecal pellets using MagMAX Microbiome Ultra Nucleic Acid Isolation kit, (Thermo Fisher Scientific, USA) and automated on KingFisher Flex robots (Thermo Fisher Scientific, USA). 16S rRNA gene amplification was performed according to the Earth Microbiome Project protocol (<u>Thompson et al. 2017</u>). Briefly, Illumina primers with unique forward primer barcodes (<u>de Muinck et al. 2017</u>) were used to amplify the V4 region of the 16S rRNA gene (515F-806R (<u>Walters et al. 2016</u>)) with single reactions per sample (<u>Marotz et al. 2019</u>). Equal volumes of each amplicon were pooled, and the library was sequenced on the Illumina MiSeq sequencing platform with paired-end 150 bp cycles.

### 16S rRNA gene data analysis

QIIME2 (<u>Bolyen et al. 2019</u>) was used to process the demultiplexed files. Sequences were filtered, denoised and trimmed to 150 bp using DADA2 (<u>Callahan et al. 2016</u>), which is used to correct Illumina-sequenced amplicon errors. The sequences were classified using the q2-feature-classifier plugin from QIIME2 that was trained on the Greengenes 13_5 (<u>McDonald et al. 2012</u>) 99% OTUs trimmed to 150 bp. [Sentence about correlation analysis.] Alpha and beta diversity plots were generated using the R microbiome package (<u>Lahti</u> <u>and Shetty 2017</u>). Alpha diversity was estimated using Faith’s phylogenetic diversity and Shannon diversity. Beta diversity was estimated using Non-Metric Multidimensional Scaling (NMDS) and Bray-Curtis dissimilarity (<u>Lahti and Shetty 2017</u>).

### *C. rodentium* induced infectious colitis

*C. rodentium* (strain DBS100) induced infectious colitis studies were performed on 8-week*-old* GIV-KO and their littermate WT control mice. *C. rodentium* were grown overnight in LB broth with shaking at 37 °C. Mice were gavaged orally with 5 x 10^8^ CFU in 0.1 ml of PBS (Bhinder et al. 2013; Koroleva et al. 2015). To determine viable bacterial numbers in faeces, fecal pellets were collected from individual mice, homogenized in ice-cold PBS, serially diluted, and plated on MacConkey agar plates. Number of CFU was determined after overnight incubation at 37 °C. Colon samples were collected to assess histology in the 7^th^ week.

### DSS-induced colitis

To induce colitis, mice were fed with drinking water containing 2.5% dextran sulfate sodium (DSS, w/v) (MP Biomedicals, MW 36–50 kDa) for five days and then replaced normal drinking water as described (<u>Chassaing</u> et al. 2014; Kiesler et al. 2015). For treatment study, MDP (100 mg/mouse/day) was administered for via intraperitoneal route in 100 ml total volume sterile saline every alternate day starting from day 0 of experiment. Mice were sacrificed on the 14^th^ day, and colon length was assessed. Colon samples were collected for assessing gene expression the levels of mRNA (by qPCR). Drinking water levels were monitored to determine the volume of water consumption. Weight loss, stool consistency, and fecal blood were recorded fro individual animals, and these parameters were used to calculate an average Disease Activity Index (DAI) as described previously (<u>Kim et al. 2012</u>). Colon histology was assessed in hematoxylin and eosin-stained tissue section using standard protocols.

### *E. coli*-induced sepsis

*E. coli* (strain RS218) was grown overnight in Lysogeny broth (LB) media with shaking at 37°C. Next morning, fresh LB media was used to dilute cultures to 1:50 and grow up to mid-log phase, washed twice with PBS, and reconstituted in PBS. GIV-KO mice (8–12-week-old) and their control WT littermates were injected with 1.5 x 10^8^ CFU of *E. coli* and mice survival was recorded for 24h post-infection. Mice were pre-treated with MDP (100 μg i.p.) or PBS 18 h before the infection.

### Transmission electron microscopy (TEM)

Cells were fixed with Glutaraldehyde in 0.1M Sodium Cacodylate Buffer, (pH 7.4) and post-fixed with 1% OsO4 in 0.1 M cacodylate buffer for 1 hr on ice. The cells were stained with 2% uranyl acetate for 1 hr on ice and dehydrated in graded series of ethanol (50-100%) while remaining on ice. The cells were washed once with 100% ethanol and twice with acetone (10 min each) and embedded with Durcupan. Sections were cut at 60 nm on a Leica UCT ultramicrotome and picked up on 300 mesh copper grids. All sections were post-stained sequentially 5min with 2% uranyl acetate and I min with Sato’s lead stain. Samples were visualised using JEOL 1400 plus equipped with a bottom-mount Gatan OneView (4k x 4k) camera.

### Protein-protein interaction network (PPIN) construction and analysis

#### Identification and construction of of NOD1, NOD2 and GIV protein protein interaction network

We have curated the interactors of NOD1 and NOD2 from a previously published BioID dataset (<u>Lu et al. 2019</u>). GIV interactors are fetched from a BioID dataset published by us (<u>Ear et al. 2021</u>) and enriched with other interactors from the human cell map (BioID repository in HEK cells (<u>Go et al. 2021</u>)). To enrich the interactions between the interactors of NOD1, NOD2 and GIV, we have fetched the high confidence edges from STRING protein interaction database (<u>Franceschini et al. 2013</u>) with a high interaction strength cutoff of 950. All such fetched interactions are combined for the protein-protein interactions network construction and visualized using Gephi. The edge list is provided in the **Supplemental Informtion 1**.

#### Topological analysis

Topological analysis was done using NetworkX library of Python. *In silico* network perturbation was performed by deletion of respective node or edge. The effect of perturbation on the protein-protein interaction network was calculated using node and edge-based centrality measurements. For each of the perturbations, changes in the centrality metrics were calculated using deference in the z score for rest of the nodes and edges to estimate the topological effect (Banerjee et al. 2015; Sinha et al. 2021).

For the node perturbation analysis GIV was deleted and the changes in the Closeness centrality and Betweenness centrality of each of the nodes in the remaining network have been calculated. Betweenness centrality, B(k) of a node k is the number of shortest paths passing through that node.

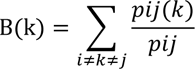

Here, pij is the number of shortest paths in the network from node I to j and pij(k) is the number of shortest paths in the network passing through k. Closeness centrality, C(k) of each node is the reciprocal of average distance of k to every other node j.

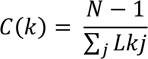

Where, N is the total number of nodes in the network and Lkj is the shortest distance between k to j.

For edge perturbation analysis the edge between GIV and NOD2 has been deleted and the change in edge betweenness and edge proximity of each of the edges in the remaining network have been calculated. Edge betweenness E of an edge, k, is the sum of the fraction of shortest paths passes through k.

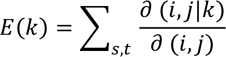

Here, `∂` (i,j) is the number of shortest paths from s to t and `∂` (i,j |k) is the number of those paths that passes through k.

Edge proximity, P, of a edge k is the closeness centrality of each edge calculated through the line graph of the actual graph.

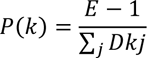

Here, E is the total number of nodes in the line graph and Dkj is the shortest distance between k to j.

For differential network analysis difference in Z score (`Δ`Z) has been calculated for each metric analysis. We take a cutoff of ΔZ >=0.5 and 1 for edge-based metrics and ΔZ >=2 for node-based metrics to identify the mostly affected edges and nodes due to GIV↔NOD2 edge and GIV deletion respectively. Identified proteins from the topological analysis are enriched to check the biological effect using CluGO. Identified nodes and edges with their difference in Z scores are provided in **Supplemental Informtion 2 and 3**.

### Plasmid constructs

The plasmids used in this study were HA-Nod1 and HA-Nod2 (from Dana Philpott (<u>Travassos et al. 2010</u>) and M. D’Amato (<u>Linderson et al. 2005</u>)); and pcDNA5-FRT-TO-RIP2-WT (from Addgene, deposited by Thomas Kufer (<u>Ellwanger et al. 2019</u>)). All other Nod2 mutants were generated by site directed mutagenesis. GST-Nod2-LRR and Myc-Nod2 were cloned into pGEX-6P and pCMV-myc vectors, respectively, using cloning sites indicated in Key Resource Table.

### Protein expression and purification

GST and His-tagged proteins were expressed in *E. coli* strain BL21 (DE3) and proteins were purified as described (Ghosh et al. 2010; Ghosh et al. 2008; Swanson et al. 2020). Briefly, to stimulate protein expression, bacteria cultures were activated with 1 mM IPTG overnight at 25°C. Bacteria were then pelleted and resuspended in either GST lysis buffer (25 mM Tris-HCL (pH 7.4), 20 mM NaCl, 1 mM EDTA, 20% (vol/vol) glycerol, 1% (vol/vol) Triton X-100, protease inhibitor cocktail) or His lysis buffer (50 mM NaH2PO4 (pH7.4), 300 mM NaCl, 10 mM imidazole, 1% (vol/vol) Triton-X-100, protease inhibitor cocktail), sonicated and lysates were cleared by centrifugation at 12,000 x g at 4°C for 30 mins. Supernatant was then affinity purified using glutathione-Sepharose 4B beads or HisPur Cobalt Resin, followed by elution, overnight dialysis in PBS, and then stored at -80°C until use.

### Transfection, lysis, and quantitative immunoblotting

HEK293T and HeLa cells were cultured in DMEM media containing 10% FBS and antibiotics according to the ATCC guidelines. Cells were transfected using polyethylenimine for DNA plasmids following the manufacturers’ protocols. Lysates for immunoprecipitation assays were prepared by resuspending cells in lysis buffer (20 mM HEPES, pH 7.2, 5 mM Mg-acetate, 125 mM K-acetate, 0.4% Triton X-100, 1 mM DTT) supplemented with 500 µM sodium orthovanadate, phosphatase inhibitor cocktails (Sigma) and protease inhibitor cocktails (Roche), and cleared (10,000 x *g* for 10 min) before use. For immunoblotting, proteins were fractionated by SDS-PAGE and transferred to PVDF membranes (Millipore). Membranes were blocked with 5% nonfat milk dissolved in PBS before incubation with primary antibodies followed by detection with secondary antibodies using infrared imaging with two-color detection and quantification were performed using a Li-Cor Odyssey imaging system. All Odyssey images were processed using Image J software (NIH) and assembled for presentation using Photoshop and Illustrator software (Adobe).

### Immunoprecipitation, *in vitro* GST-pulldown with recombinant purified protein or cell lysates

For *in vitro* pulldown assays, purified GST-tagged proteins from *E. coli* were immobilized onto glutathione Sepharose beads by incubating with binding buffer (50 mM Tris-HCl (pH 7.4), 100 mM NaCl, 0.4% (vol/vol) Nonidet P-40, 10 mM MgCl2, 5 mM EDTA, 2 mM DTT) overnight at 4ᴼC with continuous rotation. GST-protein bound beads were washed and incubated with purified His-tagged proteins resuspended in binding buffer or with pre-cleared (by centrifugation at 10,000xg for 10 min) cell lysates prepared using lysis buffer [20 mM HEPES, pH 7.2, 5 mM Mg-acetate, 125 mM K-acetate, 0.4% Triton X-100, 1 mM DTT, 500 μM sodium orthovanadate supplemented with phosphatase inhibitor cocktail (Sigma Aldrich) and protease inhibitor cocktail (Roche)] for 4 hrs at 4°C. After binding, bound complexes were washed four times with 1 ml phosphate wash buffer (4.3 mM Na2 HPO4, 1.4 mM KH2PO4 (pH 7.4), 137 mM NaCl, 2.7 mM KCl, 0.1% (vol/vol) Tween-20, 10 mM MgCl2, 5 mM EDTA, 2 mM DTT, 0.5 mM sodium orthovanadate) and eluted by boiling in Laemmli buffer (5% SDS, 156 mM Tris-Base, 25% glycerol, 0.025% bromophenol blue, 25% β-mercaptoethanol).

### Permeabilization of cells with streptolysin-O

Cells grown in 6 cm petri dishes were treated with 4 μg/ml of streptolysin-O (Sigma) for 30 min at 37°C, to permeabilize them when live to facilitate the entry of small water-soluble ATP, ATPγS or ADP nucleotides into the cells. Co-immunoprecipitation studies (described below) with or without MDP stimulation were subsequently carried out using these nucleotide-treated cells.

### Co-immunoprecipitation assays

N-terminal HA- or Myc-tagged NOD2 and C-terminal FLAG-tagged GIV full length proteins were co-expressed in HEK293T cells and 48 h after transfection, cells were stimulated with 10 µg/ml MDP (L18-MDP, Invivogen) for 1, 3 and 6 h followed by cell lysis in lysis buffer (20 mM HEPES, pH 7.2, 5 mM Mg-acetate, 125 mM K-acetate, 0.4% Triton X-100, 1 mM DTT, 0.5 mM sodium orthovanadate, Tyr phosphatase inhibitor cocktail, Ser/Thr phosphatase inhibitor cocktail, and protease inhibitor cocktail). For immunoprecipitation, equal aliquots of clarified cell lysates were incubated for 3h at 4°C with 2 μg of appropriate antibody [either anti-HA mAb or anti-FLAG M2Ab (Sigma Monoclonal ANTI-FLAG® M2, Clone M2)]. Subsequently, protein G Sepharose beads (GE Healthcare; 40 µl 50% v:v slurry) were added and incubated at 4°C for an additional 60 min. Beads were washed 4 times (1 ml volume each wash) in PBS-T buffer [4.3 mM Na2HPO4, 1.4 mM KH2PO4, pH 7.4, 137 mM NaCl, 2.7 mM KCl, 0.1% (v:v) Tween 20, 10 mM MgCl2, 5 mM EDTA, 2 mM DTT, 0.5 mM sodium orthovanadate] and immune complexes were eluted by boiling in Laemmli’s sample buffer. Bound immune complexes were separated on SDS PAGE and analyzed by immunoblotting with anti-HA, anti-Myc and anti-FLAG antibodies. In assays were the impact of nucleotides was studied on protein-protein interactions, cells were were first permeabilized with 4 μg/ml of streptolysin-O (Sigma-Aldrich, Budapest, Hungary) for 30 min at 37°C followed by incubation with nucleotides (ADP, ATP, or ATPgS).

### Confocal immunofluorescence

Cells were fixed with 4% paraformaldehyde in PBS for 30 min at room temperature, treated with 0.1 M glycine for 10 min, and subsequently blocked/permeabilized with blocking buffer (PBS containing 1% BSA and 0.1% Triton X-100) for 20 min at room temperature. Primary and secondary antibodies where incubated for 1 h at room temperature or overnight in blocking buffer. Dilutions of antibodies used were as follows: HA (1:200) and DAPI (1:1000). Alexa fluor fluorescent dye conjugated secondary antibodies were used at 1:500 dilutions. For visualizing actin organizations cells were stained with 16 mM Phalloidin Alexa Fluor^TM^-594 for 30 min at room temperature, washed three times with PBS. ProLong Glass antifade reagent is used for mounting the coverslips on glass slides.

### RAC1 activity assay

Rac1 activity in RAW 264.7 cell line was measured using GST-tagged PAK1-binding domain (PBD; pGEX-PBD) as described previously (<u>Aznar et al. 2015</u>). *E. coli* strain BL21 bacteria containing plasmid pGEX-PBD were grown at 37°C, and GST-PBD expression was induced with 1 mM IPTG (at OD600 ∼0.5) for 3 hr in 37°C incubator shaker. Lysates were prepared as described above in protein purification section, aliquoted and stored at −80°C until use. Aliquots of bacterial lysates were thawed and centrifuged to remove precipitated proteins at 14,000×g for 20 min. The clarified lysate was subsequently incubated with glutathione beads overnight at 4°C on rotating shaker to prepare bead bound GST-PBD. To analyze the role of GIV in regulation of Rac1 activity, we used RAW 264.7 cells. Cells were maintained overnight in a media containing 0.2% FBS and stimulated with or without 10 µg/ml of MDP for 0, 5 and 15 min at 37°C in CO2 incubator prior to cell lysis. Cells were lysed first with RIPA buffer [20 mM HEPES pH 7.4, 180 mM NaCl, 1 mM EDTA, 1% Triton X-100, 0.5% sodium deoxycholate, 0.1% SDS, supplemented with 1mM DTT, 500 μM sodium orthovanadate, phosphatase, and protease (Roche) inhibitor mixtures] for 15 min on ice, and then equal volume of Triton X-100 lysis buffer [20 mM HEPES, pH 7.2, 5 mM Mg-acetate, 125 mM K-acetate, 0.4% Triton X-100, 1 mM DTT, supplemented with 500 μM sodium orthovanadate, phosphatase (Sigma), and protease (Roche) inhibitor mixtures] was added for an additional 15 min. Cells were passed through a 25G needle at 4°C and lysates were subsequently cleared (10,000×*g* for 10 min). Equal volume of cleared lysates were incubated with bead-bound GST-PBD for 1 hr at 4°C with constant rotation. Beads were washed in PBS-T buffer before elution of bound proteins by boiling in Laemmli’s reduced sample buffer.

### NFkB reporter assay

RAW 264.7 cells (50000 cells/well in 96-well plate) were transfected together with 50 ng NFkB reporter plasmid and 5 ng Renilla luciferase control plasmid. After 24h, cells were stimulated with L18-MPD (10 µg/ml) for 6 hr and NFkB activity was assessed using the Dual-luciferase Reporter Assay System using manufacturers protocol. For HeLa cells 10000 cells/well in 96-well plate were seeded and transfected with 25 ng NFkB reporter plasmid, 0.5 ng Renilla luciferase control plasmid and either 5 ng/well of NOD2-WT or its mutants. Cells were primed with or without L18-MPD (10 µg/ml) for 16 hr before stimulating with LPS100ng/ml for 6h and NFkB activity was assessed using a noncommercial dual luciferase enzyme assay as described (<u>Dyer et al. 2000</u>).

### RNA extraction and Quantitative PCR

Total RNA was isolated using TRIzol reagent (Life Technologies) and Quick-RNA MiniPrep Kit (Zymo Research, USA) as per manufacturer’s guidelines. RNA was converted into cDNA using the qScript™ cDNA SuperMix (Quantabio) and quantitative RT-PCR (qPCR) was carried out using PowerUp™ SYBR™ green master mix (Applied Biosystems, USA) with the StepOnePlus Quantitative platform (Life Technologies, USA). The cycle threshold (Ct) of target genes was normalized to 18S rRNA gene and the fold change in the mRNA expression was determined using the 2^-ΔΔCt method.

### Gentamicin Protection Assay

Gentamicin protection assay was used to quantify viable intracellular bacteria as described previously (<u>Das</u> <u>et al. 2011</u>). Approximately, 2 × 10^5^ TGPMs were seeded into 12-well culture dishes overnight before infection at an MOI of 10 for 1 h in antibiotic-free RPMI media containing 10% FBS in a 37 °C CO2 incubator. Cells were then incubated with gentamicin (200 μg/ml) for 90 min after PBS was to kill extracellular bacteria. After incubation time cells were washed with PBS and subsequently lysed in 1% Triton-X 100, lysates were serially diluted and plated on LB agar plates. Bacteria colonies (CFU) were counted after overnight incubation at 37°C. To test effect of MDP, cells were pre-treated overnight with MDP (10 µg/ml).

### Cytokine Assays

Cytokines including TNFα, IL6, IL1β and IL10 were measured in cell supernatant using ELISA MAX Deluxe kits from Biolegend as per manufacturer’s protocol.

### Statistics and reproducibility

All experimental values are presented as the means of replicate experiments ±SEM. Statistical analyses were performed using Prism 9 (GraphPad Software). Differences between two groups were evaluated using Student’s t-test (parametric) or Mann-Whitney U-test (non-parametric). To compare more than three groups, one-way analysis of variance followed by Tukey’s post-hoc test was used. Differences at P < 0.05 were considered significant.

