## Supplementary Materials for "Coupling of NOD2 to GIV is Required for Bacterial Sensing"

**Affiliations:**
[1]Department of Cellular and Molecular Medicine, University of California San Diego.
[2]Department of Pathology, University of California San Diego.
[3]Department of Pediatrics, University of California San Diego.
[4]Department of Computer Science and Engineering, University of California San Diego
[5]Department of Medicine, University of California San Diego.

**Conflict of interest statement:** Authors have declared that no conflict of interest exists.

[†] Equal contribution

**KEY WORDS**: Girdin, CCDC88A, Macrophage, NOD2, MDP, Microbes, Innate immunity

***Correspondence to**:

**Pradipta Ghosh, M.D.;** Professor, Departments of Medicine, and Cell and Molecular Medicine, University of California San Diego; 9500 Gilman Drive (MC 0651), George E. Palade Bldg, Rm 232, 239; La Jolla, CA 92093. Phone: 858-822-7633: Fax: 858-822-7636:

### CATALOG OF SUPPLEMENTARY MATERIALS

1. *Supplementary Table*
2. *Supplementary Figures and Legends (S1-S6)*
3. *Supplemental Information (large excel datasheets) (1-3)*

### SUPPLEMENTARY TABLES

#### Supplementary Table 1: NOD2 mutations and its clinical relevance in CD patients.

| Mutation | Percentage of CD Patients | % CD diagnosis | Mechanistic insights | Variation Caused | Phenotypic Consequence |
|---|---|---|---|---|---|
| **1007fs (Frameshift Stop codon mutation)** | 31% | 100% | Impaired palmitoylation (Lu et al. 2019) , and protein mislocalization<br><br>Forced PM-localization of protein *does not* restore function (Lécine et al. 2007)<br><br>MDP-binding site unaffected<br><br>Loss of binding to GIV (current work) | - Mutant is unable to recognize MDP to initiate NFkB activation<br>- Associated defective release of IL-10 from blood mononuclear cells (PBMC) after stimulation with the TLR2 ligands, PGN and Pam3Cys-KKKK, and LPS. | - The genotype relative risk (GRR) for developing CD in heterozygotes and homozygotes of this mutation alone is 3.29 ±0.64 and 34.66 ±12.87 respectively<br><br>- Homozygosity is strongly associated with gastroduodenal CD and younger age at diagnosis<br><br>- Homozygous patients demonstrate a much more severe disease phenotype than other patients with Crohn's disease and have an increased risk for ileal stenoses and surgical interventions |
| **R702W (Missense Substitution Mutation)** | 32% | 100% | Impaired palmitoylation (Lu et al. 2019) , and protein mislocalization<br><br>Forced PM-localization of protein restores function (Lécine et al. 2007)<br><br>MDP-binding site unaffected<br><br>Loss of binding to GIV (current work) | - Monocyte-derived dendritic cells (MoDCs) produced significantly higher levels of IL-12 on stimulation with whole bacteria<br><br>- MoDCs carrying the mutation displayed an increased basal level of IL-8 release, which, after a bacterial encounter, equilibrated to the levels similar to healthy controls. | - GRR for developing CD in heterozygotes and homozygotes of this mutation alone is 1.97±0.85 and roughly 3.05 respectively<br><br>- Positive independent association with structuring behavior and granuloma formation |
| **G908R (Missense Substitution Mutation)** | 18% | 80% | Normal palmitoylation (Lu et al. 2019) and protein mislocalization.<br><br>Forced PM-localization of protein restores function (Lécine et al. 2007)<br><br>One of the residues that form contact site for MDP and hence, MDP recognition could be directly impacted (Vijayrajratnam et al. 2017). GIV binding (current work). | -MoDCs produced significantly higher levels of IL-12 on stimulation with whole bacteria | - GRR for developing CD in heterozygotes and homozygotes of this mutation alone is 1.97±0.85 and 4.55±1.34 respectively |

### SUPPLEMENTARY FIGURES

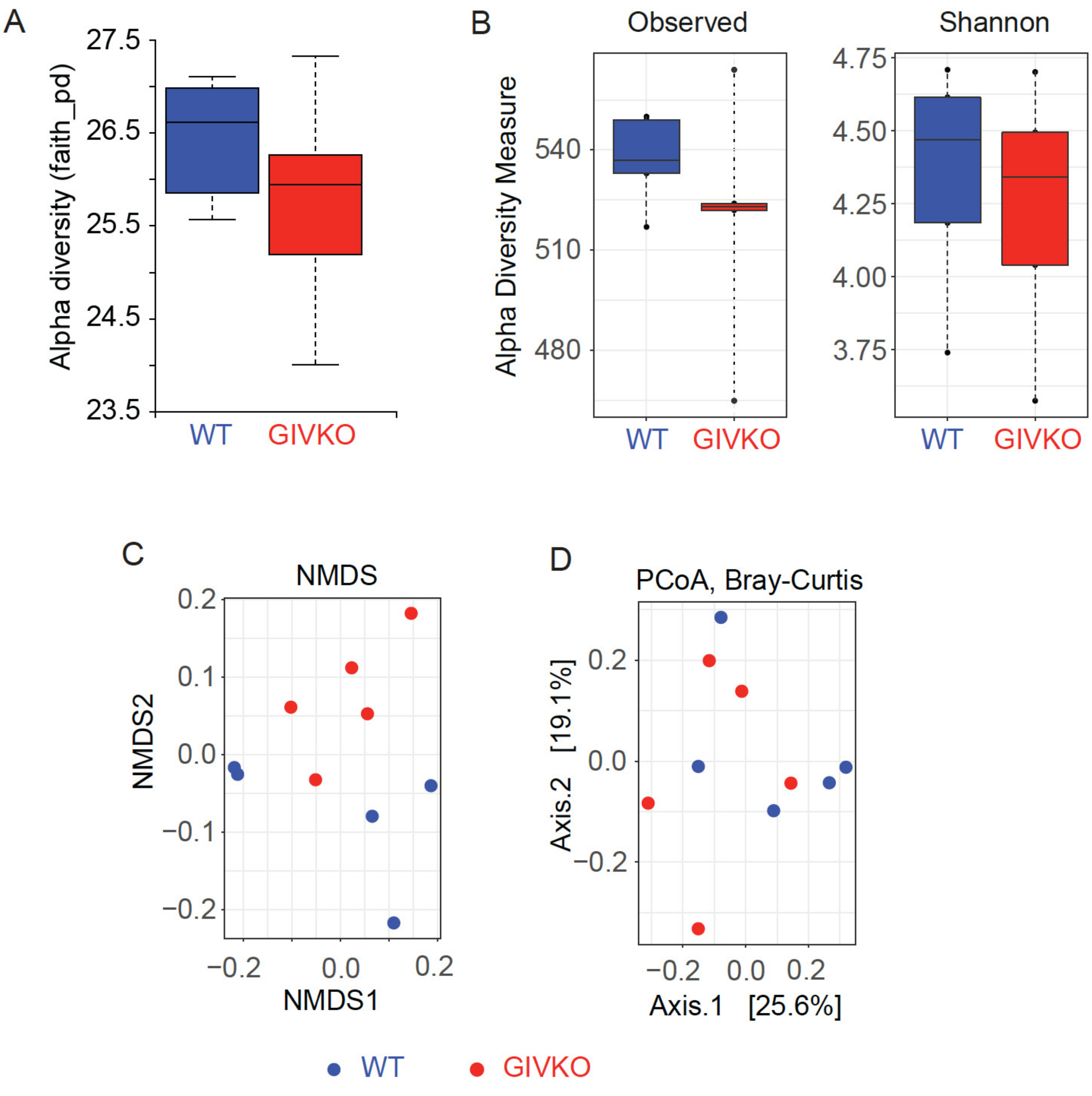


**Supplementary Figure 1:**

**Fecal microbiome analysis in 8-12 wk old mice showed that myeloid-specific (LysMCre) GIV-KO mice and develop spontaneous dysbiosis.** Related to **Figure 2A**.

**A-B**. Box plots display alpha diversity indices (Observed and Shannon), of fecal microbiome communities within GIV-KO and their littermate controls.

**C**. Non-Metric Multidimensional Scaling (NMDS) ordination of fecal microbiome communities within GIV-KO and their littermate controls

**D**. PCoA plot of beta diversity of fecal microbiome communities within GIV-KO and their littermate controls.

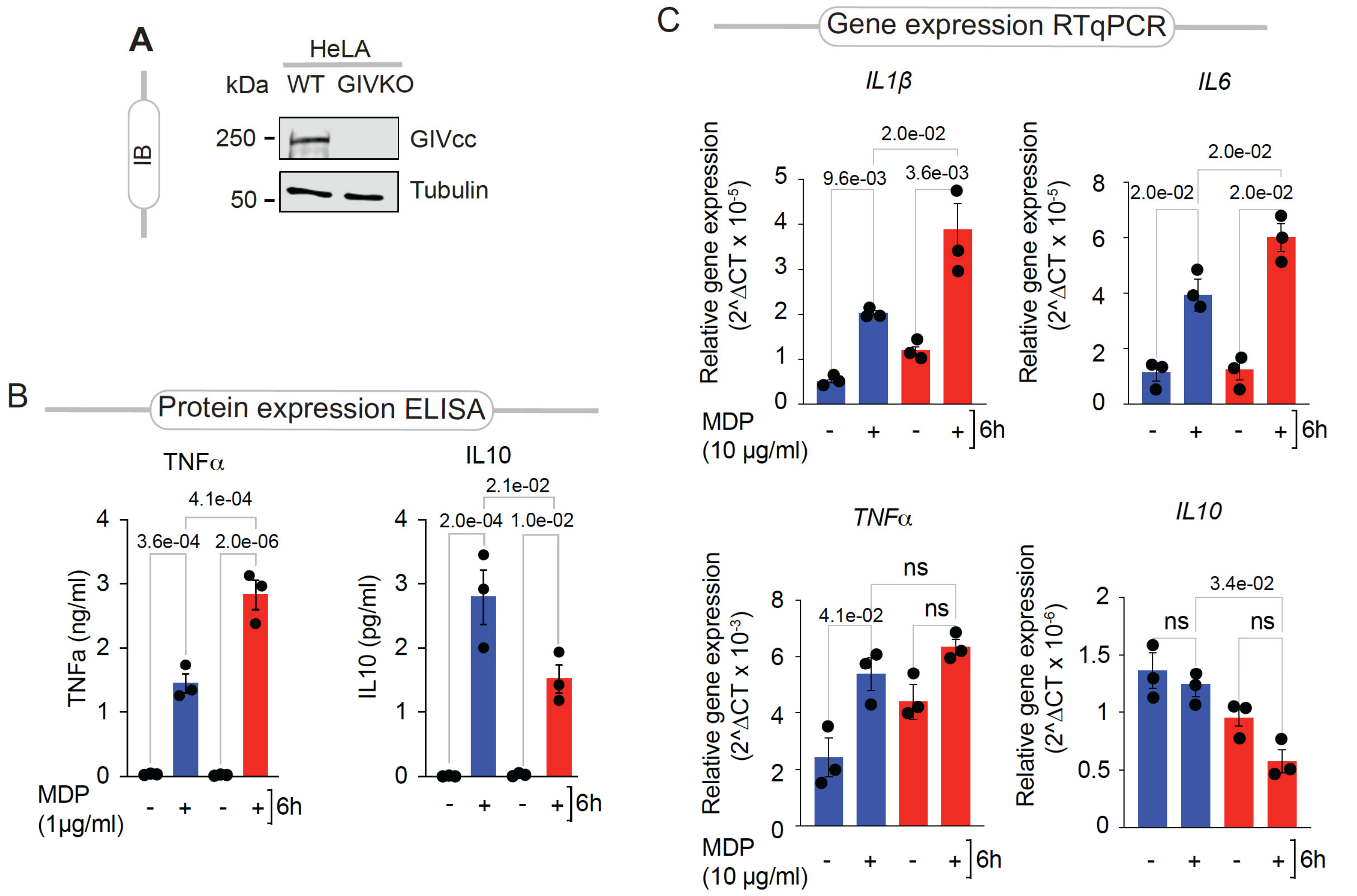


**Supplementary Figure 2: GIV is required for MDP-stimulated inflammatory resolution.** Related to **Figure 4**.

**A**. Immunoblot of control (HeLa-WT) or GIV-KO (HeLa-GIV-KO) HeLa cells.

**B.** Bar graphs display the concentrations of the indicated cytokines, as determined by ELISA.

**C.** Bar graphs display the gene expression as determined by RTqPCR. All results are displayed as mean ± SEM (n=3 biological replicates). Statistical significance was tested using two-way/one-way ANOVA followed by Tukey's test for multiple comparisons. *p*-value ≤ 0.05 is considered as significant.

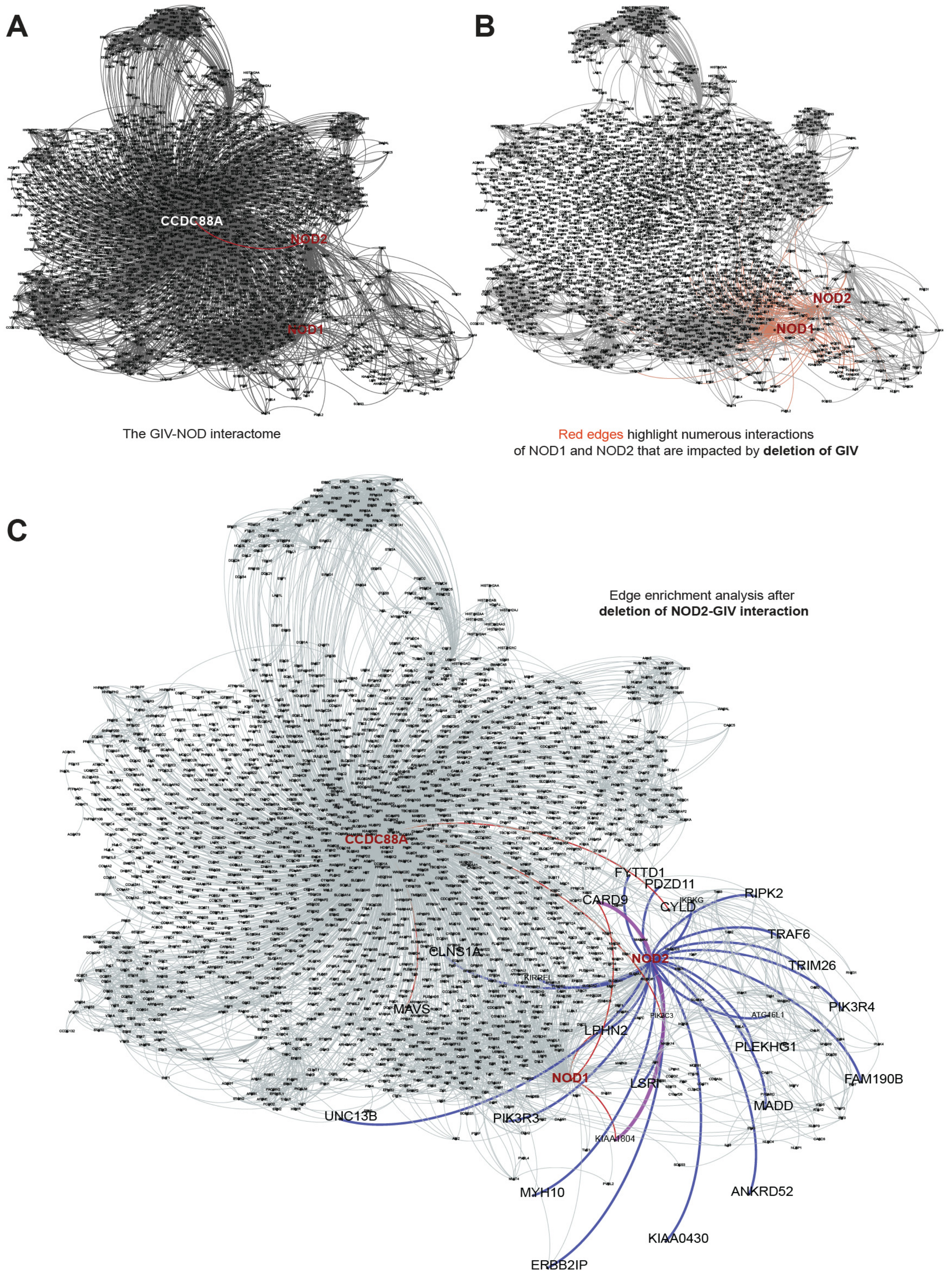


**Supplementary Figure 3: Protein-protein interaction networks, with (B-C) or without (A) *in silico* perturbation**. Related to **Figure 5A**.

**A.** The combined GIV●NOD interactome (unperturbed).

**B.** Red edges in the GIV-NOD interactome display interactions of NOD1 and NOD2 that impacted by deletion of GIV.

**C.** Highlighted (blue) interactions in GIV-NOD interactome after Edge enrichment analysis after deletion of NOD2-GIV interaction.

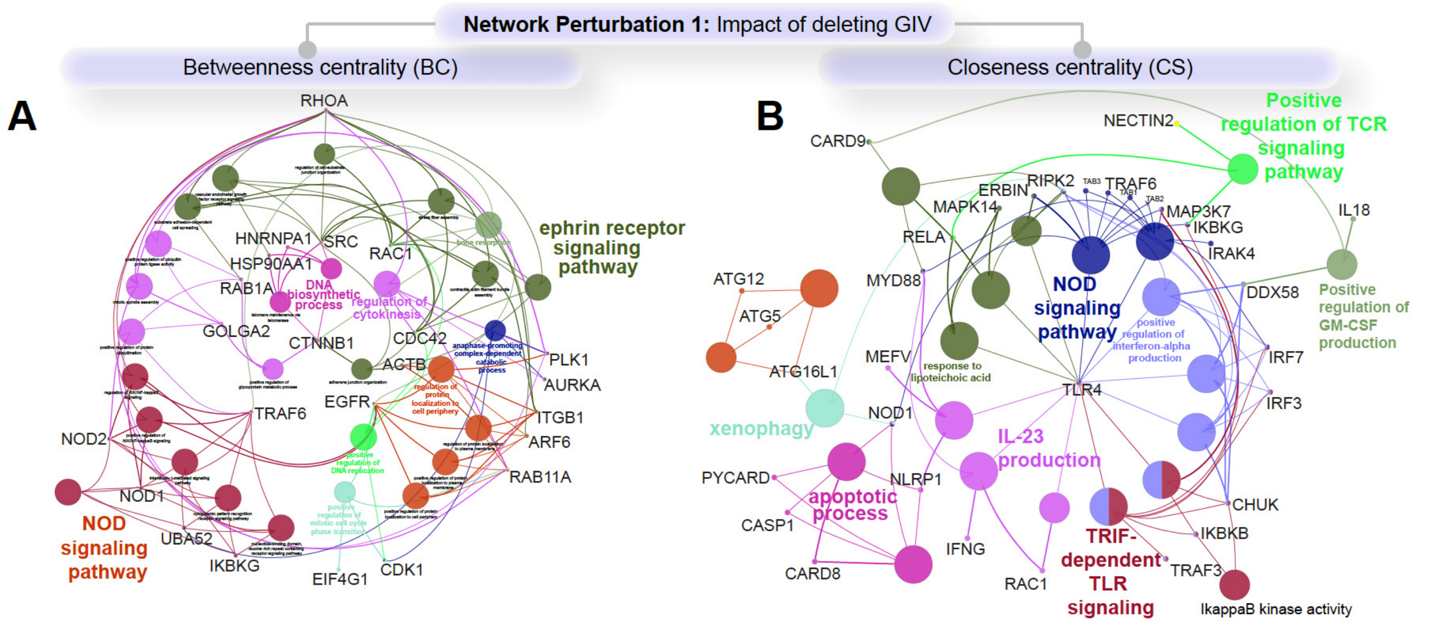


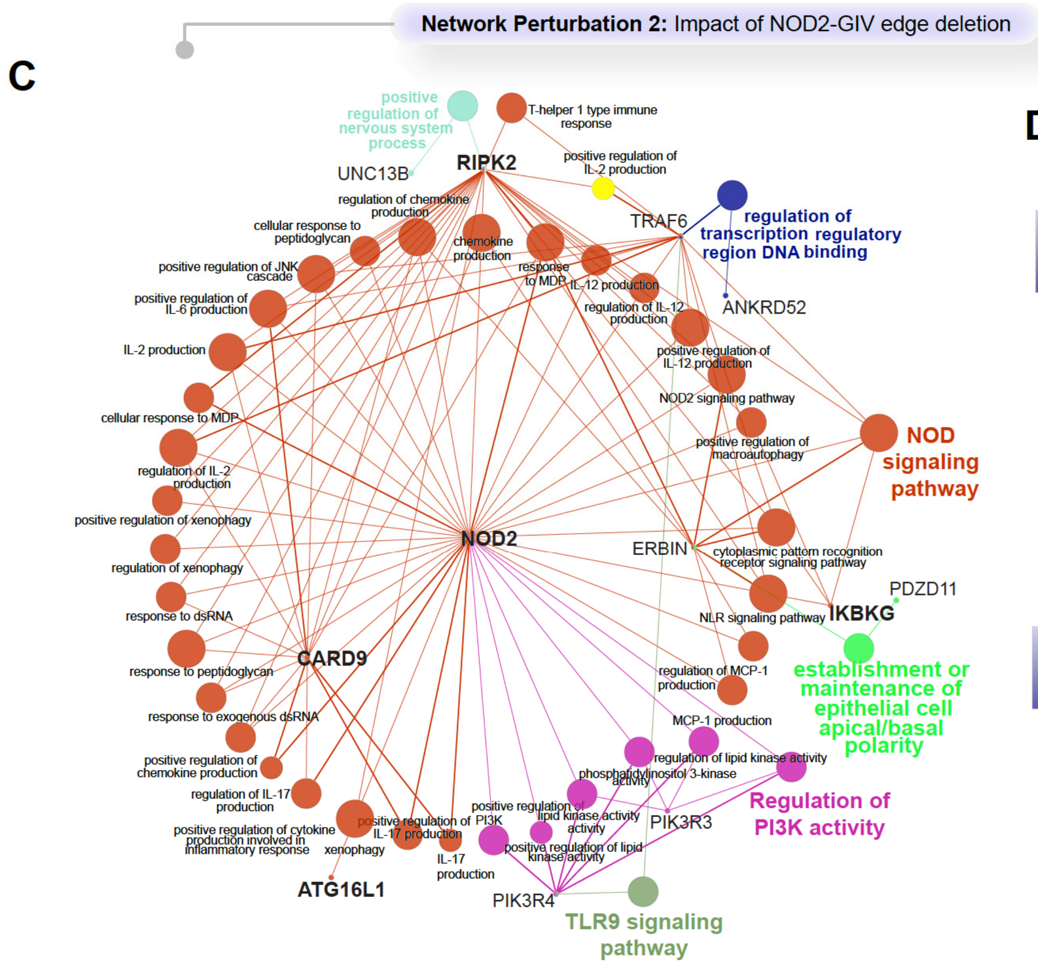


**D**

| Edge betweenness analysis | | |
|---|---|---|
| **node1** | **node2** | **Del_Z** |
| **PIK3C3** | CCDC88A | 1.8646 |
| **CCDC88A** | CYLD | 1.4692 |
| **MAVS** | CCDC88A | 1.3183 |
| **LPHN2** | NOD1 | 1.0442 |
| **KIAA1804** | NOD1 | 1.0442 |
| **CARD9** | NOD1 | 1.0349 |
| **CARD9** | NOD2 | 1.0115 |
| **KIAA1804** | NOD2 | 1.0072 |
| **LPHN2** | NOD2 | 1.0072 |

| Edge proximity analysis | | |
|---|---|---|
| **node1** | **node2** | **Del_Z** |
| **NOD2** | RIPK2 | 0.8904 |
| **NOD2** | PIK3R4 | 0.8876 |
| **ATG16L1** | NOD2 | 0.8871 |
| **NOD2** | TRIM26 | 0.8869 |
| **ANKRD52** | NOD2 | 0.8863 |
| **FAM190B** | NOD2 | 0.8863 |
| **KIAA0430** | NOD2 | 0.8863 |
| **LSRI** | NOD2 | 0.8863 |
| **MADD** | NOD2 | 0.8863 |
| **NOD2** | PLEKHG1 | 0.8863 |
| **CARD9** | NOD2 | 0.8859 |
| **LPHN2** | NOD2 | 0.8858 |
| **KIAA1804** | NOD2 | 0.8858 |
| **NOD2** | PDZD11 | 0.8857 |
| **KIRREL** | NOD2 | 0.8836 |
| **NOD2** | PIK3R3 | 0.8822 |
| **FYTTD1** | NOD2 | 0.8814 |
| **MYH10** | NOD2 | 0.8771 |
| **CLNS1A** | NOD2 | 0.8688 |
| **ERBB2IP** | NOD2 | 0.8665 |
| **NOD2** | UNC13B | 0.8546 |
| **NOD2** | TRAF6 | 0.7669 |
| **IKBKG** | NOD2 | 0.7498 |

**Supplementary Figure 4: Identification of key proteins and pathways impacted by perturbation of the NOD2●GIV network.** Related to Figure 5A.

GIV deletion in GIV-NOD interactome as perceived using betweenness Centrality (**A**) and Closeness centrality (**B**) displayed key genes and cellular process identified using CluGO analysis and Reactome pathway analysis. Impact of NOD2-GIV edge deletion identifies key genes and cellular processes (**C**) as perceived using betweenness and edge proximity analysis (**C-D**).

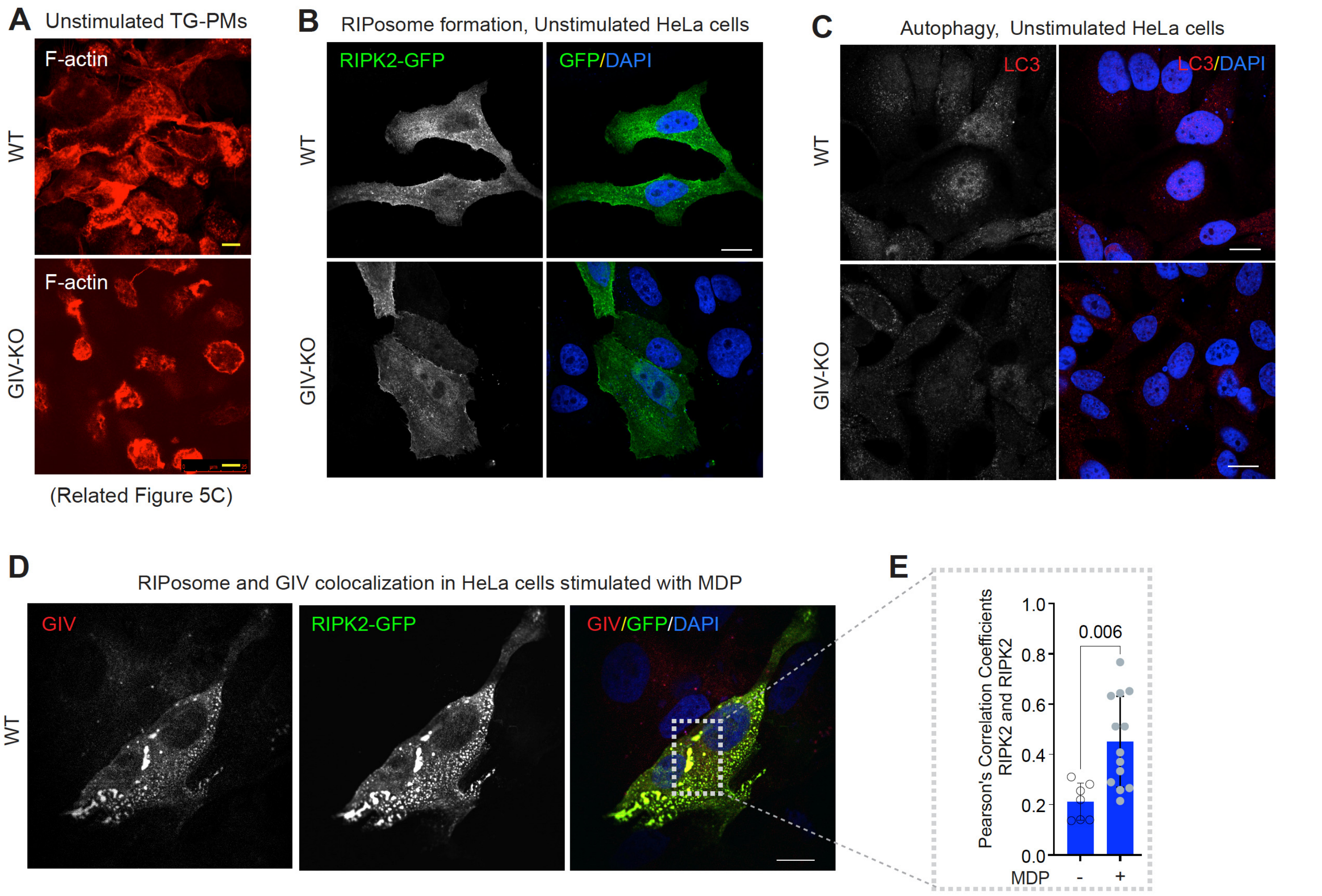


**Supplementary Figure 5: GIV and NOD2-dependent anti-bacterial cellular processes.** Related to **Figure 5**.

**A.** Representative images of unstimulated thioglycolate-stimulated murine peritoneal macrophages (TGPMs) isolated from both WT and GIVKO mice stained with phalloidin, a F-actin stain. (n = 3 mice in each group). Scale bar = 10 µm. MDP-stimulated images are displayed in Figure 5C.

**B**. Representative images of unstimulated GIV-depleted (GIV-KO) or control (WT) HeLa cells expressing GFP-RIPK2 (green = GFP; blue, DAPI/nuclei) analyzed by confocal microscopy. Scale bar = 10 µm. MDP-stimulated images are displayed in Figure 5E.

**C**. Representative images of unstimulated GIV-depleted (GIV-KO) or control (WT) HeLa cells (red = autophagosome, LC3; blue, DAPI/nuclei) analyzed by confocal microscopy. Scale bar = 10 µm. MDP-stimulated images are displayed in Figure 5F.

**D**. HeLa cells expressing GFP-RIPK2 were treated with MDP (10 µg/ml) for 6 h at 37ºC prior to fixation and staining for the markers of GIV (red), GFP (green) and DAPI (blue; nuclei) and analyzed by confocal microscopy. Colocalization analysis between GIV and RIPOsome is measured using ImageJ software. (~30-40 cells/assay; n = 3 mice in each group).

All results are displayed as mean ± SEM. Statistical significance was determined using unpaired t-test. *p*-value ≤ 0.05 is considered as significant.

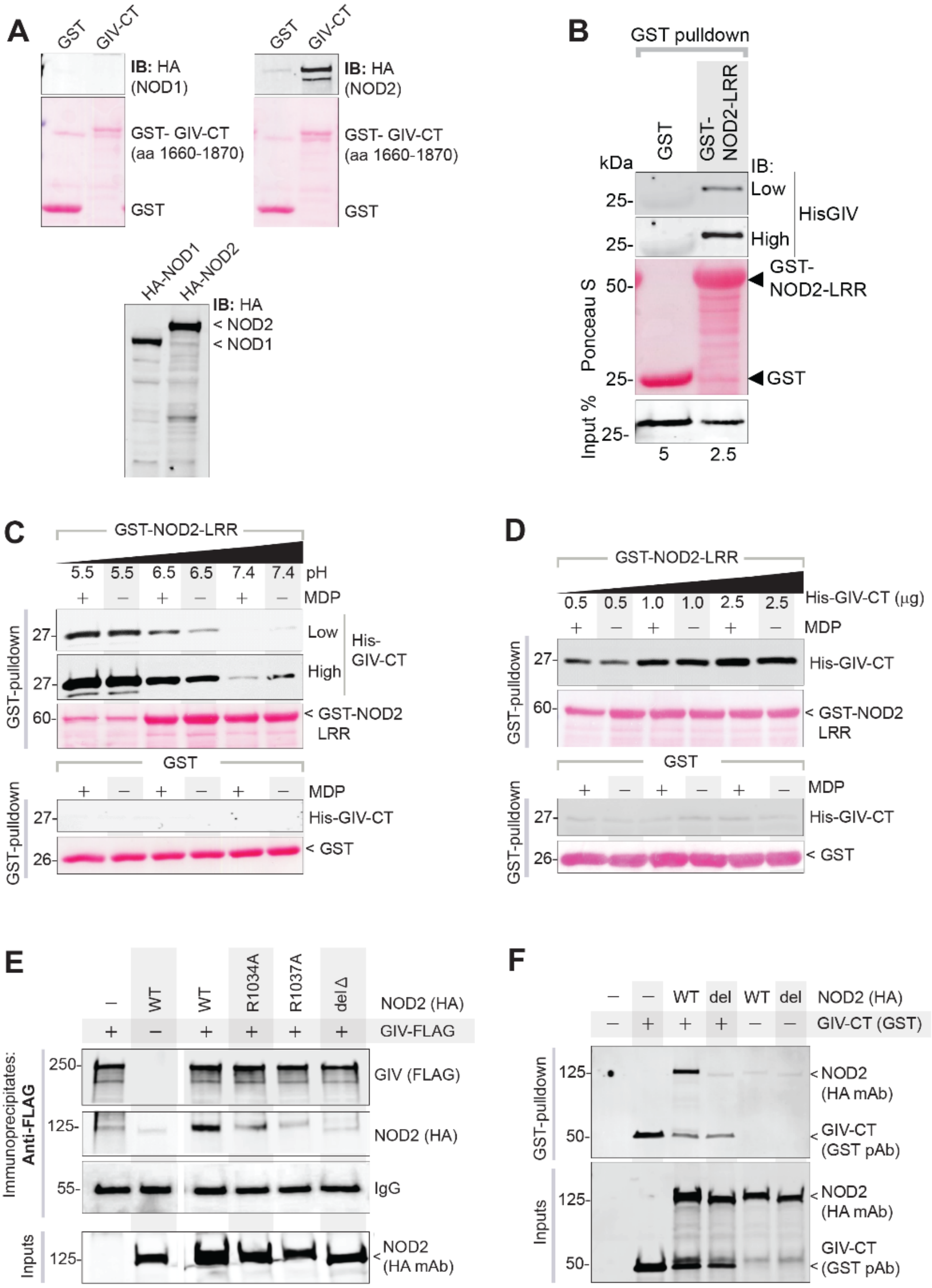


**Supplementary Figure 6: The NOD2(LRR)•GIV(C-term) interaction is a dynamic.** Related to **Figures 6-7**.
**A.** Lysates of HEK cells expressing HA-tagged NOD1/2 proteins were used in a GST pulldown assay with GST or GST-GIV-CT, and bound NOD1/2 proteins were visualized by immunoblot.

**B.** Recombinant His-GIV-CT proteins were used in a GST pulldown assay with GST or GST-NOD2-LRR, and bound GIV were visualized by immunoblot.
**C.** Recombinant His-GIV-CT proteins were used in a GST pulldown assay with GST or GST-NOD2-LRR, in the presence (+) or absence (-) of 10x molar excess of MDP at indicated pH conditions. Bound GIV was visualized by immunoblot.
**D.** Recombinant His-GIV-CT proteins (0.5, 1, 2.5 µg) were used in a GST pulldown assay with GST or GST-NOD2-LRR, in presence (+) or absence (-) of 10x molar excess of MDP, and bound GIV was visualized by immunoblot.
**E**. FLAG-tagged GIV was immunoprecipitated with anti-FLAG mAb from equal aliquots of lysates of HEK cells expressing GIV-FLAG and either wild-type (WT) or mutant HA-NOD2 constructs. Immunoprecipitated (IP; top) complexes and input (bottom) lysates were analyzed for NOD2 by immunoblotting (IB).
**F**. GST-GIV-CT was pulled down using Glutathione beads from equal aliquots of lysates of HEK lysates expressing the wild-type (WT) or del mutant HA-NOD2 construct either alone (last two lanes) or with GST-GIV-CT (aa 1660-1870; mammalian p-CEFL vector). Bound NOD2 proteins and similar expression of GIV-CT was assessed by immunoblotting (IB) using anti-HA (NOD2) and anti-GST (GIV-CT) antibodies.

### SUPPLEMENTAL INFORMATION (large excel datasheets) (1-3)

1. Supplemental information 1: The edge list (related to Figure 5A and Supplementary figure 3 and 4).
2. Supplemental information 2: Identified 'nodes' based on differences in their Z scores (related to Figure 1A and Supplementary figure 5A and 5B).
3. Supplemental information 3: Identified ‘edges’ with their differences in Z scores (related to Figure 5A and Supplementary figure 3A and 3B).
